# Asymmetric peptidoglycan editing generates the curvature of predatory bacteria, optimizing invasion and replication within a spherical prey niche

**DOI:** 10.1101/2021.06.24.449793

**Authors:** Emma J. Banks, Mauricio Valdivia-Delgado, Jacob Biboy, Amber Wilson, Ian T. Cadby, Waldemar Vollmer, Carey Lambert, Andrew L. Lovering, R. Elizabeth Sockett

**Affiliations:** Medical School, School of Life Sciences, University of Nottingham, Queen’s Medical Centre, Nottingham, NG7 2UH, UK; Institute for Microbiology and Infection, School of Biosciences, University of Birmingham, Birmingham, B15 2TT, UK; Center for Bacterial Cell Biology, Biosciences Institute, Newcastle University, Newcastle upon Tyne, NE2 4AX, UK

## Abstract

The vibrioid predatory bacterium *Bdellovibrio bacteriovorus* secretes prey wall-modifying enzymes to invade and replicate within the periplasm of Gram-negative prey bacteria. Studying self-modification of predator wall peptidoglycan during predation, we discover that Bd1075 generates self-wall curvature by exerting LD-carboxypeptidase activity in the vibrioid *B. bacteriovorus* strain HD100 as it grows inside spherical prey. Bd1075 localizes to the outer curved face of *B. bacteriovorus*, in contrast to most known shape-determinants. Asymmetric protein localization is determined by the novel function of a nuclear transport factor 2-like (NTF2) domain at the protein C-terminus. The solved structure of Bd1075 is monomeric, with key differences to other LD-carboxypeptidases. Rod-shaped Δ*bd1075* mutants invade prey more slowly than curved wild-type predators, and stretch and deform the invaded prey cell from within. Vibrioid morphology increases the evolutionary fitness of wild predatory bacteria, facilitating efficient prey invasion and intracellular growth of curved predators inside a spherical prey niche.

## Introduction

*Bdellovibrio bacteriovorus* HD100 is a small, vibrioid-shaped predatory bacterium which invades and then replicates within the periplasm of Gram-negative prey bacteria, forming a spherical structure called a prey bdelloplast^1^. *B. bacteriovorus* has a broad prey range which includes multidrug-resistant pathogens with variable outer membrane and cell wall chemistries, and occurrence of genetic resistance to *B. bacteriovorus* has never been observed in prey bacteria^2, 3^. Predatory *B. bacteriovorus* can also successfully clear pathogen infections within a range of *in vivo* animal models^4,5,6^ and therefore has considerable and growing potential as a novel antimicrobial therapeutic.

The predation process is critically dependent upon the modification of both predator and prey peptidoglycan (PG) cell walls to facilitate the dual bacterial encounter. PG forms a complex macromolecular structure called a sacculus which surrounds the cytoplasmic membrane of nearly all bacteria, maintaining cell shape and providing protection against lysis due to osmotic pressure fluctuations and large extracellular toxins^7^. Bacterial growth, cell division, and – importantly in this study – predation, occur through PG remodeling which involves a repertoire of predator-secreted modifying enzymes^8,9,10,11^.

The predatory lifecycle of *B. bacteriovorus* begins with attack-phase cells which swim^12^ or glide^13^ to encounter prey, then recognize and attach to the prey outer membrane. An entry porthole in the prey cell wall is created, through which the predator traverses to enter the inner periplasmic compartment^10^. Concurrently, two predator DD-endopeptidases are secreted into prey, cleaving cross-links between prey PG peptide chains to sculpt rod-shaped prey cells into spherical bdelloplasts^8^. This also reduces the frequency of sequential predator invasions, thus conferring exclusivity to the first-entering predator^8^. The porthole in the wall and outer membrane is then re-sealed and the predator secretes hydrolytic enzymes including nucleases and proteases into the cytoplasm of the now-dead host, taking up the nutrient-rich degradative products^14, 15^. Prey-derived and *de novo*-synthesized nucleotides are incorporated into the replicating genome copies of the predator, which grows as an elongating multi-nucleoid filament inside the rounded but intact prey until exhaustion of prey nutrients^16^. Synchronous septation of the predator filament yields progeny cells which secrete targeted PG hydrolytic enzymes to lyse the prey host and re-initiate the predatory cycle^11^.

PG hydrolases have an additional role generally in the determination of cell shape^17^, which has been particularly studied in the non-predatory, ε-proteobacteria *Helicobacter pylori*^18,19,20^ and *Campylobacter jejuni*^21,22,23^, in whom multiple PG hydrolases collectively generate helical morphology. In contrast, bacterial vibrioid morphology is generally determined by non-enzymatic cyto- or periskeletal proteins (well-studied in *Caulobacter crescentus*^24, 25^ and *Vibrio cholerae*^26^).

Despite the characterization of predator enzymes which modify the prey PG, there have been very few studies concerning the cell wall PG architecture or vibrioid cell shape of predatory bacteria. Here, we investigate the mechanism by which a curved, vibrioid predator is generated and ask whether there are evolutionary and functional connections between predator cell morphology and an efficient predatory lifestyle.

We identify and characterize the first predatory cell shape-determinant: Bd1075, which is targeted to the outer convex cell face by its C-terminal nuclear transport factor 2-like (NTF2) domain, where it exerts localized LD-carboxypeptidase (LD-CPase) activity upon the *B. bacteriovorus* PG wall to generate curvature and the classical vibrio shape. The Bd1075 protein has some novel features in comparison to other LD-CPases, being monomeric with a C-terminal extension to the NTF2 domain binding pocket. We discover that rod-shaped Δ*bd1075* mutant predators invade prey more slowly than the curved wild-type and stretch and deform the prey cell bdelloplast while growing within, unlike the curved wild-type. We further note that there is dynamic adaptation to the spherical prey niche; both curved wild-type and rod-shaped Δ*bd1075* mutant predators temporarily adopt a curve while growing inside the spherical prey bdelloplast, however only wild-type predators exit prey with a permanent vibrioid shape.

Our findings suggest that the evolution of a vibrioid cell shape confers two fitness advantages to *B. bacteriovorus* predators: rapid prey entry and optimal replication within a spherical intra-bacterial niche. This discovery also implies a possible scenario in which cell curvature may first be “templated” by predatory growth inside a spherically-shaped structure, then sensed and permanently “fixed” by PG shape-determining enzymes.

## Results

### Bd1075 generates the curvature of *B. bacteriovorus* predators

The monocistronic *bd1075* gene of vibrioid-shaped *B. bacteriovorus* Type strain HD100 encodes a 329 amino acid hypothetical protein with a predicted N-terminal sec signal peptide^27^, suggestive of protein translocation into the periplasm or secretion from the cell (Supplementary Fig. 1). Bd1075 shares limited homology with Csd6 (identity: 24%, similarity: 38%) and Pgp2 (identity: 25%, similarity: 40%) which are dimeric proteins important for the generation of helical cell shape in *H. pylori*^18, 28^ and *C. jejuni*^21^, respectively. These comparisons led us to hypothesize that Bd1075 could fulfill a role in the shape-determination of vibrioid predator *B. bacteriovorus*.

Reverse-transcriptase PCR (RT-PCR) revealed that *bd1075* is constitutively transcribed throughout the predatory cycle, suggesting that the protein may have a role in *B. bacteriovorus* rather than a secreted predatory function (Supplementary Fig. 2).

A markerless deletion of *bd1075* in the curved *B. bacteriovorus* Type strain HD100 could still be cultured predatorily (phenotype differences further detailed later) but Δ*bd1075* mutant cells had a distinct straight rod-shaped morphology unlike the curved wild-type HD100 parent strain (Fig. 1a-b). Wild-type mean curvature was significantly higher than the Δ*bd1075* mutant (0.61 A.U. ± SD 0.33 *versus* 0.17 A.U. ± SD 0.22, respectively, p<0.0001; Fig. 1c). Plasmid-based complementation of Δ*bd1075* with the wild-type *bd1075*_HD100_ gene increased curvature relative to the Δ*bd1075* mutant (Fig. 1c). These results indicate that Bd1075 has a role in generating the curvature of *B. bacteriovorus*.

**Fig. 1.**
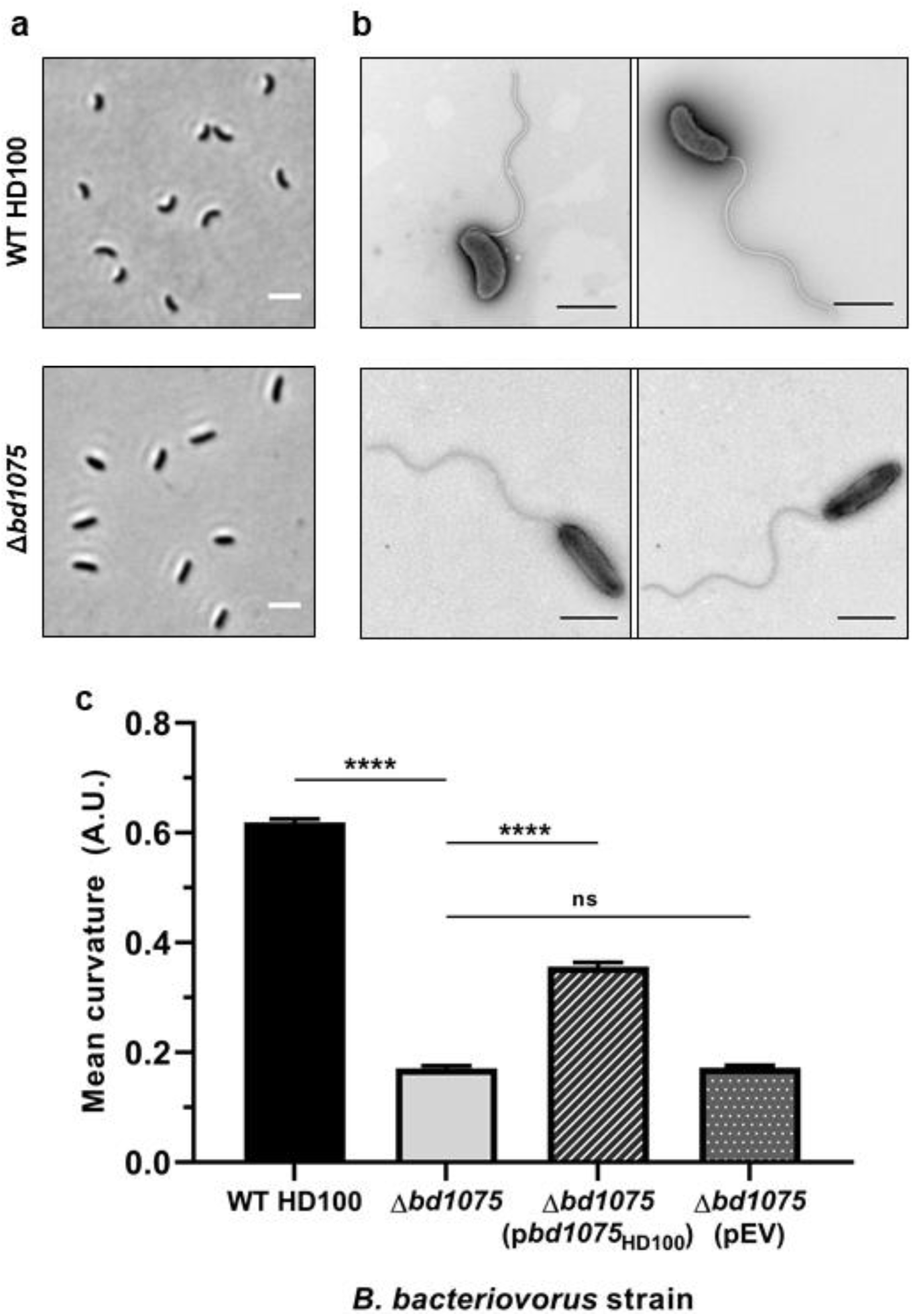
Bd1075 generates the curvature of *B. bacteriovorus* HD100 predator cells. **a** Phase-contrast images of attack-phase *B. bacteriovorus* cells showing the curvature of wild-type (WT) HD100 cells in comparison to non-vibrioid Δ*bd1075* cells. Images are representative of cells from at least 5 biological repeats. Scale bars = 2 µm. **b** Transmission electron micrographs of WT HD100 and Δ*bd1075* cells stained with 0.5% uranyl acetate. Scale bars = 1 µm. Images are representative of 3 biological repeats. **c** Curvature measurements of *B. bacteriovorus* attack-phase cells. n = 1920-2503 cells per strain from 3 biological repeats. Error bars represent standard error of the mean. ns: non-significant (p>0.05), ****: p<0.0001; Kruskal-Wallis test. Frequency distributions are included in Supplementary Fig. 5a.

### A lab-evolved predator strain that cannot generate cell curvature

The straight rod morphology of Δ*bd1075* resembles the long-cultured laboratory strain *B. bacteriovorus* 109J which was isolated in the 1960s and is the only reported non-vibrioid strain of *B. bacteriovorus*^1^. As *bd1075* is conserved in all *B. bacteriovorus* strains including 109J and appears to be a curvature-determinant, we queried why strain 109J is non-vibrioid. Despite otherwise 100% sequence identity with *bd1075*_HD100_, *bd1075*_109J_ contains an in-frame N-terminal truncation of 57 amino acids (Pro-18 to Tyr-74) (Supplementary Fig. 3a-c). RT-PCR confirmed that *bd1075*_109J_ is expressed in attack-phase cells and that the RNA transcript contains the predicted truncation (Supplementary Fig 3d). To test whether the N-terminal truncation may render the translated protein non-functional in curvature-determination, we cross-expressed *bd1075*_109J_ in the HD100 Δ*bd1075* mutant and this did not complement curvature (Supplementary Fig. 4). In contrast, cross-expression of *bd1075*_HD100_ in wild-type 109J significantly increased the curvature of strain 109J (0.34 A.U. ± SD 0.26 *versus* 0.22 A.U. ± SD 0.21, respectively, p<0.0001; Supplementary Fig 4)

These results show that Bd1075 is the curvature-determinant of vibrioid *B. bacteriovorus* strains, and that an inactivating mutation within the gene resulted in the lab-evolved strain 109J which is unable to generate cell curvature.

### Rod-shaped Δ*bd1075* predators invade prey more slowly than the curved wild-type

As cell morphology can be phenotypically important in other bacteria, we asked whether the curved shape of *B. bacteriovorus* could be advantageous to the bacterial predator during its unique intraperiplasmic lifecycle. Comparison of the gross predation efficiency of wild-type and Δ*bd1075* predators upon *E. coli* prey in either liquid culture, or on pre-grown *E. coli* biofilms, did not reveal a significant difference (Supplementary Fig. 6 and Supplementary Fig. 7). However, these are laboratory conditions with readily-available prey and in which multiple important factors required to locate and navigate towards prey (e.g. predator chemotaxis and locomotion) are operational in bringing predators close to the prey surface.

We considered that predator morphology may fulfill an important role at the interface of single predator-prey encounters and therefore studied predation more closely at the single-cell level using time-lapse microscopy to visualize individual predatory invasion events. *B. bacteriovorus* HD100 wild-type or Δ*bd1075* strains were mixed with *E. coli* S17-1 and placed under a microscope which captured images of specific fields of view every 1 min until the majority of *E. coli* prey had been invaded. Hypothesizing that curvature may affect the invasion of *B. bacteriovorus* into prey, we measured two parameters: prey attachment time and prey entry time (Fig. 2a). Duration of prey attachment did not significantly differ (p = 0.46) between the wild-type (29.3 min ± SD 5.0) and Δ*bd1075* (29.6 min ± SD 4.8) (Fig. 2b), however there was a significant difference (p<0.0001) between the rates at which wild-type and Δ*bd1075* entered prey: 4.3 min ± SD 0.9 *versus* 6.1 min ± SD 1.7, respectively (Fig. 2c). Moreover, the longest wild-type entry was a single 7 min invasion, whereas 35.6% of Δ*bd1075* entry invasions were ≥ 7 min, with the longest invasion lasting 14 min.

**Fig. 2.**
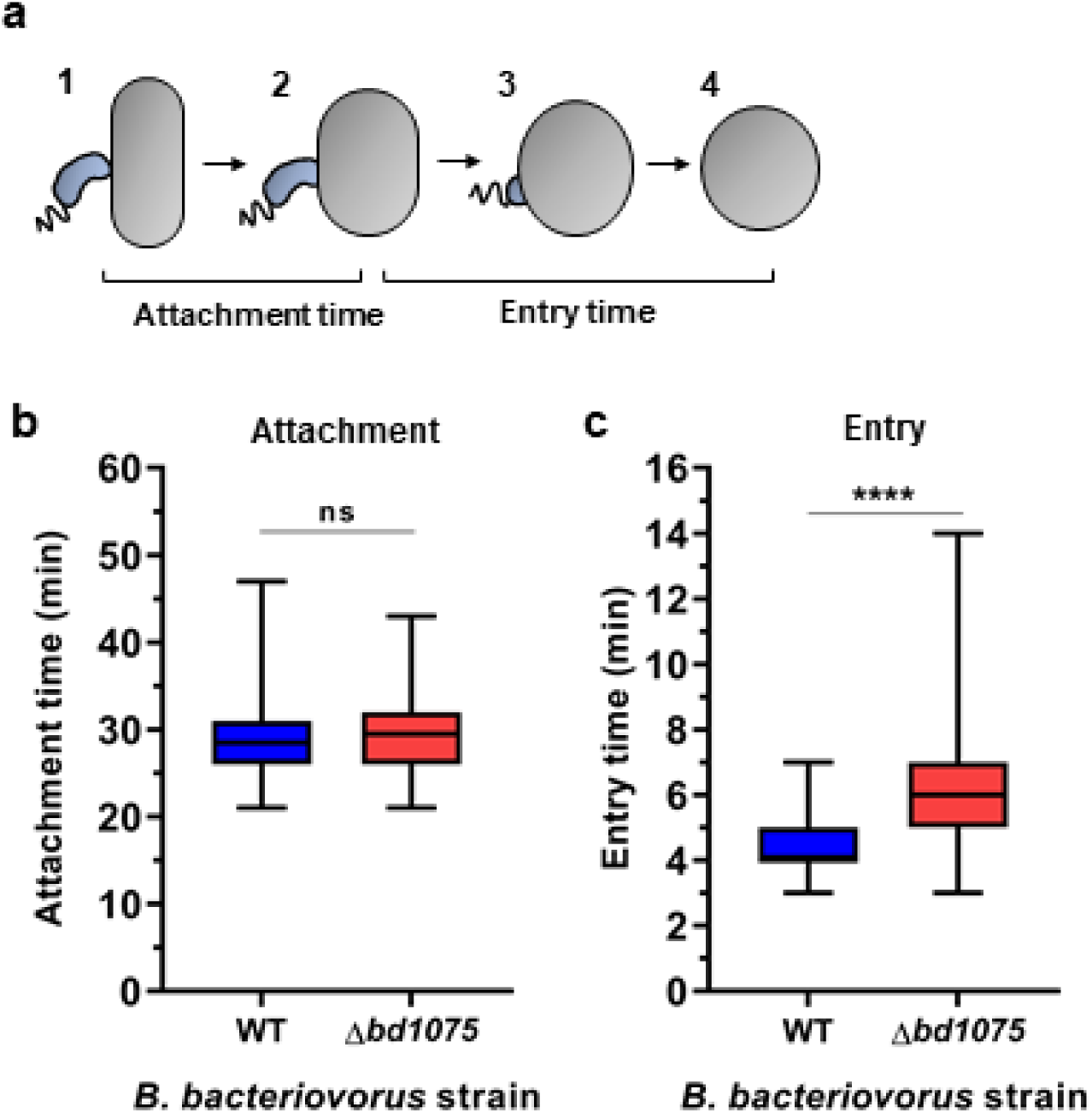
Prey attachment and entry times of *B. bacteriovorus* wild-type and Δ*bd1075*. **a** Schematic to illustrate the measurement of attachment and entry times. Attachment time: number of frames (1 frame = 1 min) between initial predator attachment to prey and the first sign of predator entry into prey (stages 1-2). Entry time: number of frames between the first sign of predator entry and the predator residing completely inside the prey bdelloplast (stages 2-4). **b** Duration of attachment to and **c** entry into *E. coli* S17-1 prey by *B. bacteriovorus* HD100 wild-type (WT) and Δ*bd1075*, measured by time-lapse microscopy. n = 90 cells in total from 3 biological repeats. Box: 25^th^ to 75^th^ percentiles; whiskers: range min-max; box line: median; ns: non-significant (p>0.05); ****: p<0.0001; Mann-Whitney test.

These data indicate that *B. bacteriovorus* vibrioid morphology facilitates the traversal of predators across the prey cell envelope into the intraperiplasmic compartment of the rounded prey cell.

### Prey bdelloplasts are stretched and deformed by replicating, non-curved predators

Having observed that the non-vibrioid Δ*bd1075* mutant was slower to enter the prey periplasm than the curved wild-type, we next investigated the growth and replication of Δ*bd1075* within prey. During prey invasion, *B. bacteriovorus* secretes two DD-endopeptidases, Bd0816 and Bd3459, into the prey periplasm which hydrolyze the peptide bonds connecting chains of polysaccharide backbone^8^. The prey PG wall becomes more malleable and the cell rounds up into a spherical bdelloplast.

We hypothesized that growth of the straight rod-shaped Δ*bd1075* within spherical bdelloplasts may be deleterious to the predatory niche, whereas curved wild-type cells may better “fit” into the curvature of the bdelloplast during growth and elongation. A C-terminal mCerulean3 fusion to the continuously-expressed cytoplasmic protein Bd0064^6, 29^ was introduced via single-crossover recombination into both wild-type HD100 and Δ*bd1075* to label the predator cytoplasm blue and allow visualization of *B. bacteriovorus* within prey. Fluorescent *B. bacteriovorus* strains were mixed with *E. coli* S17-1 pZMR100 and observed throughout the predatory cycle. Wild-type predators elongated as tightly curved filaments inside bdelloplasts (Fig. 3a), however the rod-shaped Δ*bd1075* mutant - despite becoming more curved by the spherical bdelloplast environment over time (Fig. 3b) - elongated as a less tightly curved filament and then septated to give rod-shaped, non-vibrioid progeny cells (Fig. 3a). Strikingly, a sub-set of Δ*bd1075* predator cells appeared to stretch and deform the usually spherical prey bdelloplasts during intra-bacterial growth (Fig. 3c).

**Fig. 3.**
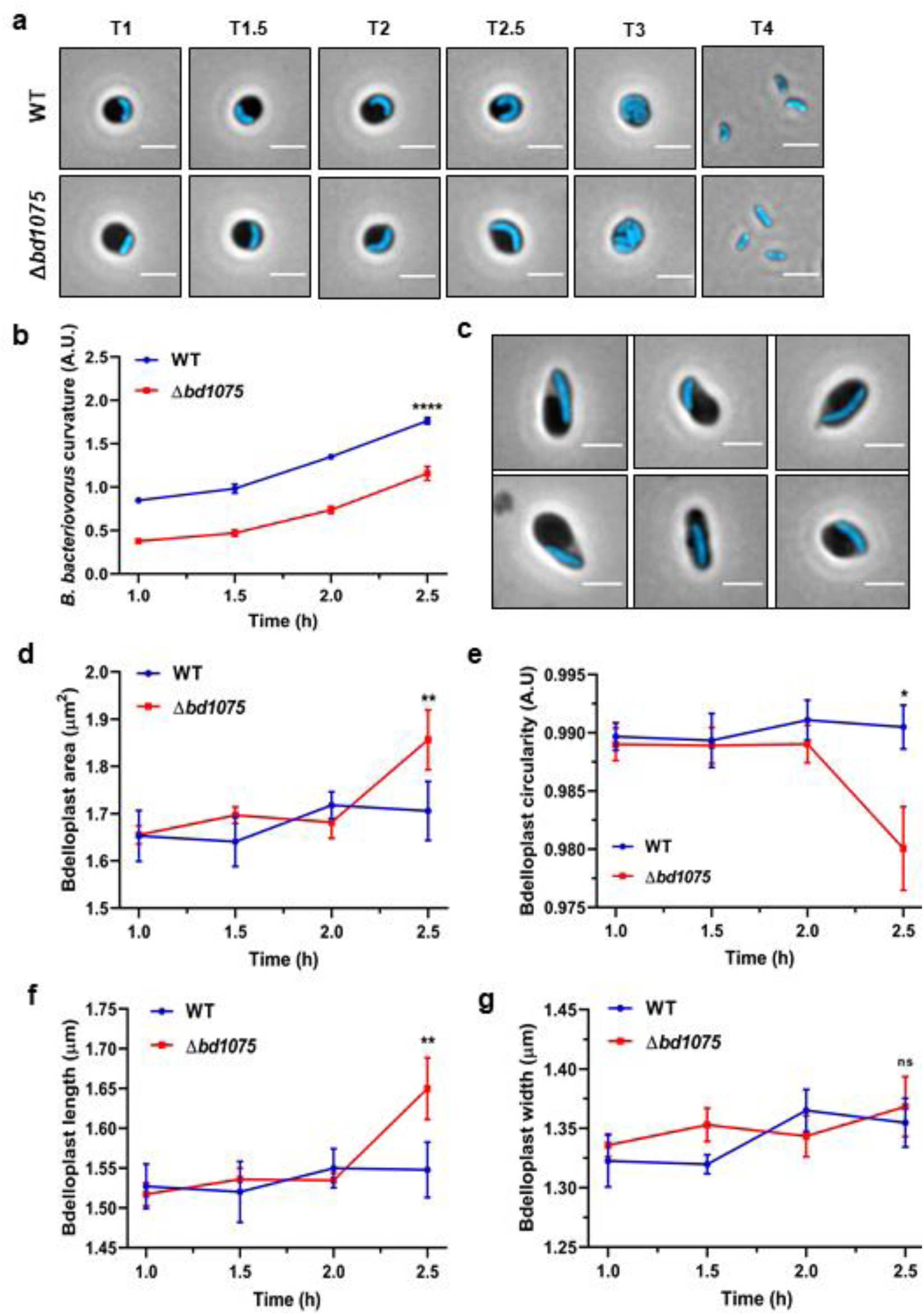
Intra-bacterial growth and bdelloplast topology effects of *B. bacteriovorus* strains. **a** Growth of *B. bacteriovorus* wild-type (WT) and Δ*bd1075* strains inside *E. coli* S17-1 pZMR100 prey bdelloplasts. *B. bacteriovorus* strains express the cytoplasmic fusion protein Bd0064-mCerulean3 to allow visualization of intraperiplasmic predator cells. T = hours elapsed since predators and prey were mixed. Scale bars = 2 µm. Images are representatives of cells from 3 biological repeats. **b** Curvature of *B. bacteriovorus* WT and Δ*bd1075* strains during predation upon *E. coli* S17-1 pZMR100 as depicted in (**a**). n = 134-250 cells per strain and per timepoint from 3 biological repeats. Error bars represent standard error of the mean. ****: p<0.0001; Mann-Whitney test. **c** Examples of Δ*bd1075* cells which appear to stretch and deform the *E. coli* prey bdelloplast at T = 2.5 h during 3 repeats of predatory timecourses as shown in (**a**). Scale bars = 2 µm. **d** Area, **e** circularity, **f** length and **g** width of *E. coli* prey bdelloplasts during predation by WT or Δ*bd1075* predators. n = 134-250 cells per strain and per timepoint from 3 biological repeats. Error bars represent standard error of the mean. ns: non-significant (p>0.05); **: p<0.01, *: p<0.05; Mann-Whitney test.

Measuring the morphology of bdelloplasts containing a single *B. bacteriovorus* predator between 1 h and 2.5 h after predator-prey mixing showed that from 1 h - 2 h, the shape of prey bdelloplasts did not obviously differ between the two strains. However, at 2.5 h when *B. bacteriovorus* cells are nearing maximal growth, the morphology of prey bdelloplasts became markedly different. The area of each bdelloplast containing a Δ*bd1075* mutant predator (1.87 μm^2^ ± SD 0.57) was significantly higher than bdelloplasts containing wild-type predators (1.69 μm^2^ ± SD 0.32, p<0.01; Fig. 3d) and the circularity was significantly lower (Δ*bd1075:* 0.98 A.U. ± SD 0.04; wild-type: 0.99 A.U. ± SD 0.01, p<0.05; Fig. 3e). Bdelloplasts containing Δ*bd1075* predators were also significantly longer (1.65 μm ± SD 0.36, p<0.01; Fig. 3f) than those containing curved wild-type predators (1.54 μm ± SD 0.18) but the width did not significantly differ (Δ*bd1075:* 1.38 μm ± SD 0.13; wild-type: 1.35 μm ± SD 0.09, p>0.05; Fig. 3g), consistent with the visual appearance of “stretched” bdelloplasts.

Collectively, these findings show that curved *B. bacteriovorus* wild-type cells have an “optimal fit” into the curvature of the bdelloplast, whereas the shape of rod-shaped predators can result in severe deformation of the spherical prey niche.

### Bd1075 demonstrates LD-carboxypeptidase activity on PG sacculi *in vivo* and *in vitro*

Bd1075 contains a predicted LD-transpeptidase (LDT) catalytic domain (Supplementary Fig. 1c-d), however the LDT domains of related proteins Csd6 and Pgp2 function instead as LD-carboxypeptidases (LD-CPases) which remove the terminal D-alanine from a PG tetrapeptide (consisting of L-Ala, D-Glu, *meso*-Dap, and D-Ala) to generate a tripeptide (consisting of L-Ala, D-Glu, and *meso*-Dap)^30^. This highlighted the need to verify the catalytic activity (if any) of Bd1075. PG sacculi were therefore purified from *B. bacteriovorus* strains and analyzed by HPLC to determine their muropeptide composition and any changes to it caused by Bd1075.

In contrast to curved wild-type HD100 sacculi, rod-shaped Δ*bd1075* sacculi contained a greater proportion of monomeric tetrapeptides (23.7% ± 0.8%) and cross-linked tetratetrapeptides (33.2% ± 0.7%) compared to the wild-type (9.6% ± 0.8% and 18.6% ± 0.6%, respectively), and a complete absence of monomeric tripeptides and dimeric tetratripeptides (Fig. 4a-b and Table 1). This difference suggests that Bd1075 could cleave the C-terminal D-alanine of tetrapeptides to produce tripeptides which terminate with *meso*-Dap. The complemented strain Δ*bd1075* (p*bd1075*_HD100_) contained no monomeric tetrapeptides, 14.8% ± 1.2% tripeptides and 7.2% ± 0.5% dipeptides. These data suggest that re-introduction of the wild-type *bd1075*_HD100_ gene resulted in over-complementation beyond wild-type as all monomeric tetrapeptides have been cleaved to tripeptides, with some subsequently converted to dipeptides (Fig. 4c and Table 1).

**Table 1.** Quantification of muropeptides released from B. bacteriovorus HD100 sacculi. Values represent the relative percentage area of each muropeptide peak in **Fig. 4**. Numbers with an asterisk differ from WT HD100 by more than 30% and values that are additionally emboldened differ by more than 50%.

| Muropeptide | Relative peak area (%)<sup>1</sup> in *B. bacteriovorus* strain |  |  |  |
| --- | --- | --- | --- | --- |
|  | WT HD100 | *Δbd1075* | *Δbd1075*<br>(*pbd1075*<sub>HD100</sub>) | *Δbd1075*<br>(pEV) |
| **Monomers** |  |  |  |  |
| Tri | 14.4 ± 1.0 | **n.d.\*<sup>2</sup>** | 14.8 ± 1.2 | **n.d.\*** |
| Tetra | 9.6 ± 0.8 | **23.7 ± 0.8\*** | **0.6 ± 0.9\*** | **23.1 ± 2.7\*** |
| Di | 2.1 ± 0.1 | **n.d.\*** | **7.2 ± 0.5\*** | **n.d.\*** |
| Penta | 12.8 ± 2.5 | 11.9 ± 3.6 | 12.2 ± 4.2 | 14.8 ± 0.0 |
| Monomer anhydroMurNAc | 2.2 ± 3.0 | 2.3 ± 3.2 | 2.6 ± 3.7 | 2.5 ± 3.6 |
| Monomers (Total) | 38.9 ± 0.5 | 35.6 ± 4.3 | 34.9 ± 2.6 | 37.8 ± 2.6 |
| **Dimers** |  |  |  |  |
| TetraTri | 14.9 ± 0.7 | **0.5 ± 0.7\*** | **26.4 ± 1.5\*** | **n.d.\*** |
| TetraTetra | 18.6 ± 0.6 | **33.2 ± 0.7\*** | **7.8 ± 1.4\*** | **29.6 ± 4.7\*** |
| TetraPenta | 12.2 ± 1.0 | 12.5 ± 0.3 | 14.7 ± 2.9 | 13.1 ± 0.2 |
| Dimer anhydroMurNAc | 17.3 ± 0.8 | 16.4 ± 0.6 | 20.5 ± 4.4 | 17.0 ± 0.3 |
| Dimers (total) | 45.6 ± 2.3 | 46.1 ± 0.6 | 48.9 ± 2.9 | 42.7 ± 4.5 |
| **Trimers** |  |  |  |  |
| TetraTetraTri | 1.6 ± 0.4 | **n.d.\*** | **3.8 ± 0.2\*** | **n.d.\*** |
| TetraTetraTetra | 6.9 ± 0.5 | **11.2 ± 0.8\*** | 4.9 ± 1.0 | **10.6 ± 2.2\*** |
| TetraTetraPenta | 6.9 ± 3.6 | 7.1 ± 3.2 | 7.4 ± 4.3 | 8.8 ± 4.0 |
| Trimer anhydroMurNAc | 7.0 ± 0.3 | 9.5 ± 1.0* | 6.2 ± 1.3 | 10.3 ± 1.0* |
| Trimers (total) | 15.4 ± 2.8 | 18.3 ± 4.0 | 16.2 ± 5.5 | 19.4 ± 1.8 |
| Dipeptides (total) | 2.1 ± 0.1 | **n.d.\*** | **7.2 ± 0.5\*** | **n.d.\*** |
| Tripeptides (total) | 22.4 ± 0.6 | **0.2 ± 0.3\*** | 29.3 ± 0.5* | **n.d.\*** |
| Tetrapeptides (total) | 54.3 ± 1.1 | 79.3 ± 2.0* | 41.4 ± 4.2 | 75.7 ± 1.4* |
| Pentapeptides (total) | 19.0 ± 1.3 | 18.2 ± 0.9 | 19.5 ± 0.6 | 21.8 ± 5.0 |
| AnhydroMurNAc (total) | 13.1 ± 3.3 | 13.6 ± 2.6 | 14.9 ± 5.4 | 14.5 ± 3.8 |
| Average chain length | 7.9 ± 2.0 | 7.5 ± 1.4 | 7.2 ± 2.6 | 7.1 ± 1.9 |
| Degree of cross-linkage | 33.1 ± 0.7 | 35.3 ± 2.8 | 35.3 ± 2.2 | 34.3 ± 1.0 |
| % peptides in cross-links | 61.1 ± 0.5 | 64.4 ± 4.3 | 65.1 ± 2.6 | 62.2 ± 2.6 |

<sup>1</sup> values are mean ± variation of two biological replicates.
<sup>2</sup> n.d., not detected.

**Fig. 4.**
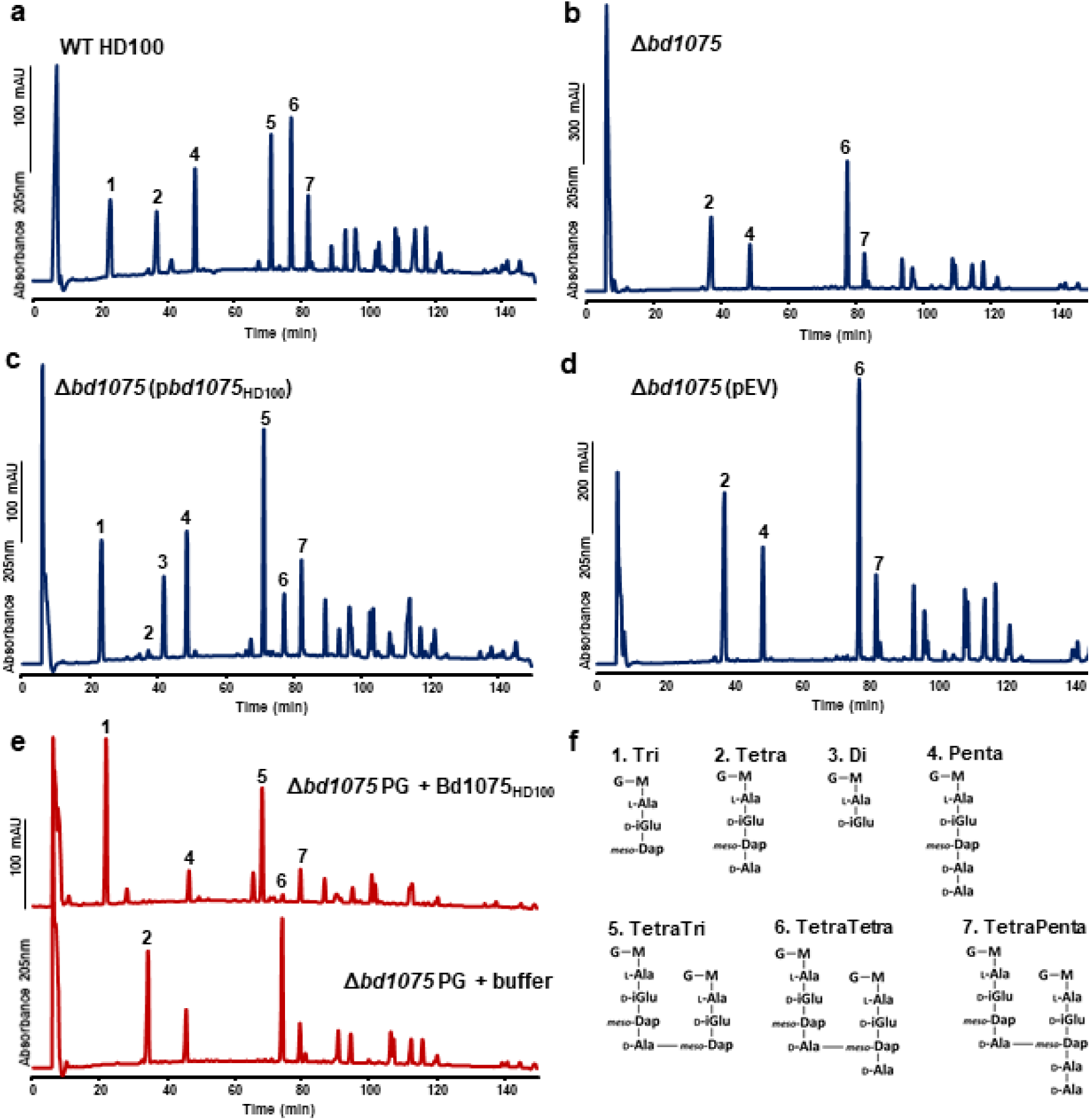
Muropeptide composition of *B. bacteriovorus* HD100. **a-d** HPLC elution profiles of reduced muropeptides released from peptidoglycan sacculi isolated from attack-phase *B. bacteriovorus* HD100 cells. **a** Wild-type (WT) HD100, **b** Δ*bd1075*, **c** Δ*bd1075* (p*bd1075*HD100) - *bd1075*HD100 expressed in Δ*bd1075*, and **d** Δ*bd1075* (pEV) - empty vector control in Δ*bd1075*. Representative chromatograms of 2 biological repeats are shown. **e** HPLC muropeptide elution profiles of Δ*bd1075* sacculi treated with either purified Bd1075HD100 enzyme (above) or buffer control (below). Data are from 1 biological repeat. **f** Structural schematics of the seven primary muropeptide fractions. Numbers correspond to those above peaks in (**a-e**). G: *N*-acetylglucosamine, M: *N*-acetylmuramitol, L-Ala: L-alanine, D-Glu: D-glutamic acid, *meso*-Dap: *meso*-diaminopimelic acid, D-Ala: D-alanine. Minor peaks are annotated in Supplementary Fig. 10 (for **a-d**) or Supplementary Fig. 9a (for **e**).

In comparison, the muropeptide profile of strain Δ*bd1075* (p*bd1075*_109J_), in which curvature was not complemented (Supplementary Fig. 4), did not differ from Δ*bd1075*, further confirming the non-functionality of truncated Bd1075_109J_ as an LD-CPase (Supplementary Fig. 8a and Supplementary Table 1).

Wild-type 109J had a very similar muropeptide profile to the Δ*bd1075* mutant - a complete absence of tripeptides and tetratripeptides and a high proportion of monomeric tetrapeptides (28.0% ± 4.3%) and dimeric tetratetrapeptides (33.8% ± 1.1%) (Supplementary Fig. 8b and Supplementary Table 1). Finally, the cross-complementation strain 109J (p*bd1075*_HD100_), with an increased mean curvature compared to the lab-cultured wild-type strain 109J (Supplementary Fig. 4), contained a higher proportion of monomeric tripeptides (18.7% ± 3.0%) and dimeric tetratripeptides (26.2% ± 0.6%), and a reduction in monomeric tetrapeptides and dimeric tetratetrapeptides (3.0% ± 0.8% and 8.8% ± 0.8%, respectively) (Supplementary Fig. 8c and Supplementary Table 1). This demonstrates that cross-expression of *bd1075*_HD100_ in wild-type 109J resulted in enzymatic conversion of tetrapeptides to tripeptides and increased the curvature of this normally non-vibrioid strain.

To further validate the LD-CPase activity of Bd1075, an N-terminally His-tagged copy of *bd1075*_HD100_ was expressed in *E. coli* BL21 and purified to near homogeneity by Ni-NTA affinity chromatography and size-exclusion chromatography. The muropeptide profile of *B. bacteriovorus* HD100 Δ*bd1075* sacculi incubated with purified Bd1075 enzyme revealed a complete conversion of both monomeric tetrapeptides to tripeptides and dimeric tetratetrapeptides to tetratripeptides (Fig. 4e). Bd1075 had identical enzymatic activity on wild-type 109J straight rod sacculi and upon sacculi of wild-type *E. coli* BW25113, showing that the enzyme can act on PG from different bacterial strains and species (Supplementary Fig. 9b-c).

These muropeptide data determine that Bd1075 has LD-CPase activity on PG both *in vivo* and *in vitro*, removing C-terminal D-alanine residues linked to the L-center of *meso*-Dap to convert tetrapeptides to tripeptides.

### Bd1075 structure determination

The structure of mature Bd1075 protein was determined to 1.34 Å (Fig. 5 and Supplementary Table 2). The Bd1075 structure contains two domains: the catalytic LD-CPase domain (aa 47-180) and a C-terminal nuclear transport factor 2 (NTF2)-like domain (aa 196-304) (Fig. 5a). Interestingly, although there was a global agreement in fold to Csd6 of *H. pylori* and Pgp2 of *C. jejuni* which also contain an NTF2 domain and an LD-CPase domain (the junction between the two at residue 188 of Bd1075), there were significant differences in fold elements and local regions. These differences resulted in an inability to solve Bd1075 via molecular replacement, necessitating the use of SAD phasing with co-crystallized halide ions. The Bd1075 protein is monomeric (the two molecules in the asymmetric unit contact one another by packing interactions only), lacking the dimerization regions of the other characterized LD-CPase proteins; this was supported by size exclusion data (Supplementary Fig. 11).

**Fig 5.**
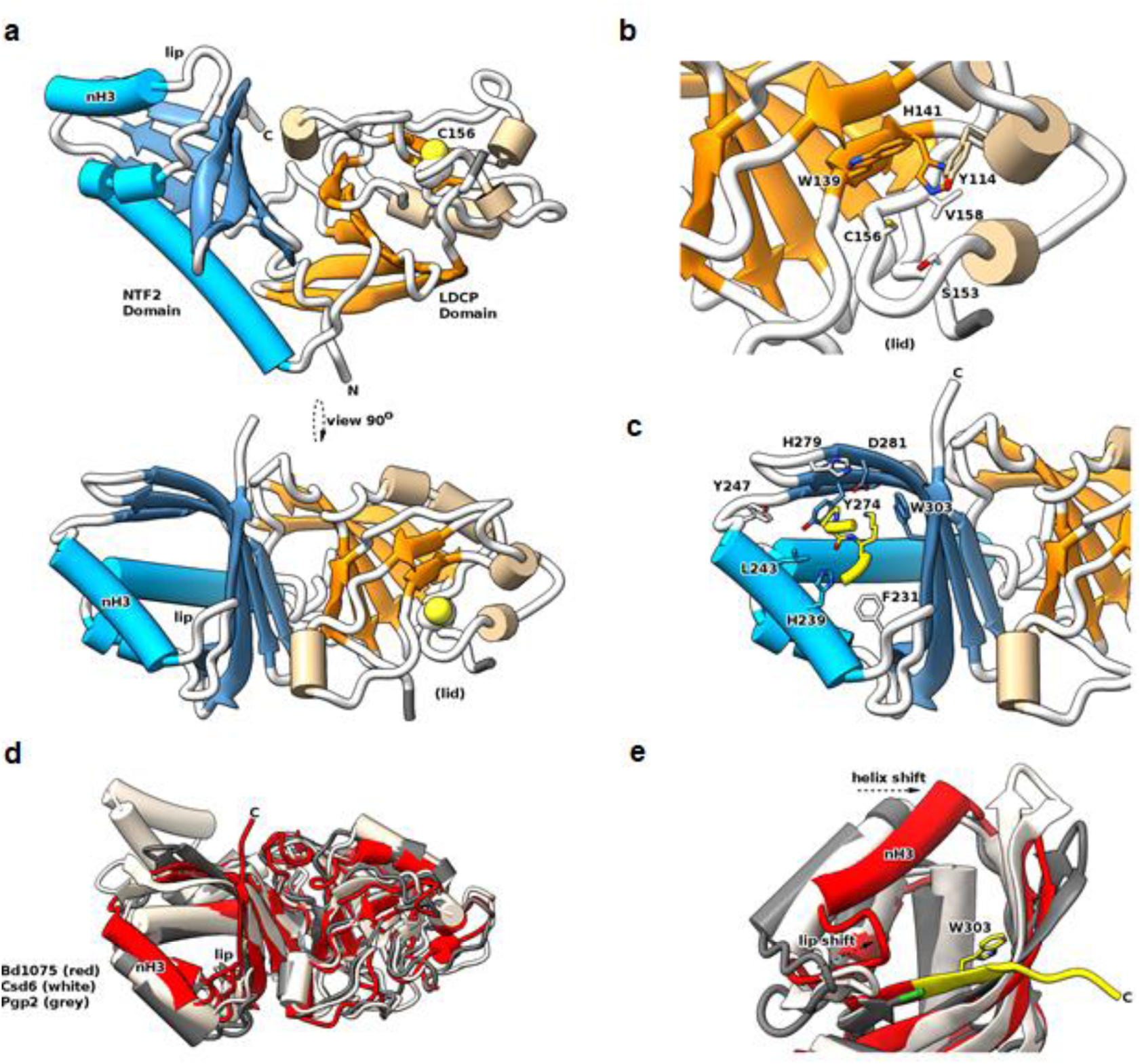
Structure of Bd1075 and features different to other characterized LD-CPase enzymes. **a** Two orthogonal views of the Bd1075 fold, with catalytic residue C156 in space-fill form and features labeled. **b** Close-up view of the Bd1075 LD-CPase catalytic domain with selected residues that form the active site pocket displayed in stick form. **c** Close-up view of the Bd1075 NTF2 pocket, demonstrating complexation of a loop (residues 106-109 colored yellow, P107 and K108 in stick form) from a neighboring molecule in the crystal lattice. **d** Comparison of Bd1075 (red, 7O21), Csd6 (white, 4XZZ) and Pgp2 (gray, 6XJ6) structures. Helix 3 of the Bd1075 NTF2 domain (labeled “nH3”) and the associated loop (“lip”) are relatively closer to the NTF2 pocket than the respective features of Csd6/Pgp2. **e** Close-up of the NTF2 terminus from structural alignment in (**d**), demonstrating the relative extension of the Bd1075 C-terminus (colored yellow, includes NTF2 pocket-forming residue W303) in comparison to the shorter Csd6/Pgp2 termini (end residue colored green). The relative shifts of the nH3 helix and lip loop to constrict the NTF2 pocket are denoted by dashed arrows.

We were able to trace residues 29-308 with the exception of a presumably flexible region (aa 82-91) which we term the active site “lid”. Differences to other LD-CPase structures are distributed throughout the fold (and in a small shift in NTF2:LD-CPase juxtaposition) as demonstrated by RMSD values for the full-length/LD-CPase-alone/NTF2-alone of 2.5 / 2 / 1.8 Å for Csd6 and 2.8 / 1.9 / 2.2 Å for Pgp2. The large values of 2.5/2.8 Å for full-length RMSD are in contrast with the agreement of 1.8 Å between Csd6 and Pgp2, hence Bd1075 is the structural outlier of the three proteins.

Bd1075 has a consensus active site, with the superfamily conserved catalytic triad consisting of C156, H141 and A142, each present in the expected (presumed active) orientations (Fig. 5b). A142 is often a glycine residue in other LD-CPases but here it makes identical h-bonding contacts to H141 using backbone carbonyl atoms. Bd1075 active site pocket-forming residue V158 is a relative anomaly as this position is an arginine in most LD-transpeptidases or an alanine in Csd6/Pgp2 and YafK-like enzymes shown recently to cleave the cross-link between PG and Braun’s lipoprotein^31, 32^. The termini of Bd1075 are very different to both Csd6 and Pgp2, replacing the N-terminal dimerization domain of those enzymes with a simple, shorter loop, and extending the C-terminus such that the Bd1075 NTF2 domain finishes with a longer beta-strand (aa 295-306) - residues of which contribute to its binding pocket (Fig. 5d-e). The C-terminus has a further 21 residues that we were unable to fit in this crystal form and which are predicted to be at least partly disordered.

Most noteworthy within the core of the Bd1075 C-terminus was the presence of W303 which is clearly located within the substrate binding pocket of the NTF2 domain. W303 is highly conserved amongst *Bdellovibrio* Bd1075 protein homologs but not found in other LD-CPase proteins (due to their shorter NTF2 sequences which terminate at a position equivalent to Bd1075^E302^). The crystal packing of Bd1075 was such that a two amino acid loop of P107/K108 from one Bd1075 monomer packed into the NTF2 domain of the adjacent monomer. This loop is situated in an identical position to the bound glycerol of Csd6, postulated to be reminiscent of a substrate-like interaction^28^. Conserved residues of the Bd1075 NTF2 domain binding pocket pack around this feature (yellow in Fig. 5c), the base of which is formed by Y274 – an important residue in both our monomeric structure and other dimeric LD-CPases.

Having begun to probe the cellular localization of Bd1075 in *B. bacteriovorus,* we used this new structural information to aid in the construction of fluorescently-tagged Bd1075 truncations and point mutants for enzyme localization tests.

### The C-terminal NTF2 domain of Bd1075 targets the protein to what becomes the outer convex cell face, and is required to generate curvature

#### Localization and targeting of Bd1075 to the outer convex cell face

To determine whether Bd1075 is broadly active over all *B. bacteriovorus* envelope PG or if activity is specifically localized, a double-crossover markerless strain in which mCherry is C-terminally fused to Bd1075 was constructed. Bd1075-mCherry localized to the outer convex face of *B. bacteriovorus* - both in free-swimming attack-phase cells (Fig. 6a) and throughout the predatory cycle (Supplementary Fig. 12). This contrasts with most characterized vibrioid cell shape-determinants in other bacteria such as *Caulobacter crescentus* and *Vibrio cholerae*, which typically localize to the inner concave face^24, 26^.

**Fig. 6.**
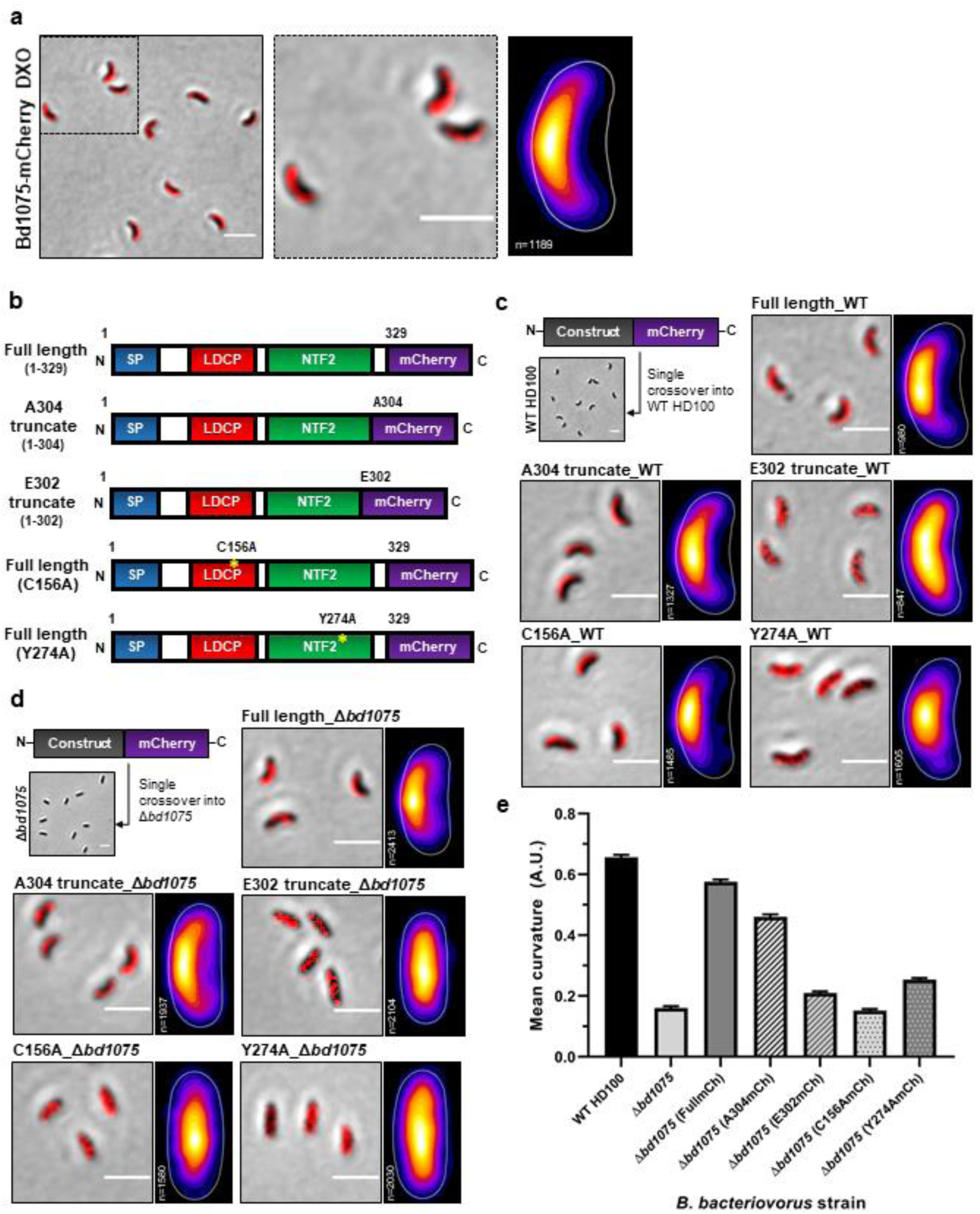
The NTF2 domain is required to target Bd1075 to what becomes the convex cell face and generate curvature at that site. **a** *B. bacteriovorus* Bd1075-mCherry double-crossover (DXO) attack-phase cells (left), showing the localization of wild-type Bd1075-mCherry to the convex cell face, and representative of 3 biological repeats. Dashed boxed region is shown in a close-up (middle). Scale bars = 2 μm. Heatmap (right) depicts the location of wild-type Bd1075-mCherry foci detected in n = 1189 cells from 3 biological repeats. White-yellow = highest intensity, purple-black = lowest intensity. **b** Schematics of Bd1075-mCherry single-crossover constructs used in (**c-e**). Full-length: Residues 1-329 (wild-type complete protein) fused to mCherry, A304: Residues 1-304 (contains a completed NTF2 domain including the *B. bacteriovorus*-specific residue W303) fused to mCherry, E302: Residues 1-302 (does not complete the NTF2 domain) fused to mCherry, C156A: Residues 1-329 (full-length with a point mutation of C156A in the catalytic LD-CPase domain) fused to mCherry, and Y274A: Residues 1-329 (full-length with a point mutation of Y274A in the NTF2 domain) fused to mCherry. **c-d** Bd1075-mCherry single-crossover constructs introduced into either *B. bacteriovorus* HD100 **c** wild-type (contains a native wild-type copy of *bd1075*) or **d** Δ*bd1075* (lacking a wild-type copy of *bd1075*). Attack-phase cell images and adjacent heatmaps show targeting of Bd1075-mCherry. Images and heatmaps were generated from 3 biological repeats (n = number of cells analyzed). Scale bars = 2 μm. **e** Curvature measurements of *B. bacteriovorus* Δ*bd1075* attack-phase cells containing different single-crossover Bd1075-mCherry fusions. n = 1886-2812 cells per strain from 3 biological repeats. Error bars represent standard error of the mean. All pairwise comparisons between strains (except for Δ*bd1075 vs* Δ*bd1075* (C156mCh) were significant (p<0.0001; Kruskal-Wallis test). Frequency distributions are included in Supplementary Fig. 5c.

Moreover, unlike pole-associated proteins, the mechanism by which bacterial proteins are targeted to one of the cell lateral sides has not been described. Intrigued and partly informed by the protein structural data, we utilized the specifically-targeted Bd1075 to investigate how bacterial proteins might be targeted to just one lateral wall. We hypothesized that, in addition to the N-terminal signal peptide which targets the protein for translocation into the periplasm, Bd1075 may contain a second internal targeting sequence which directs the protein to just one side of the periplasm: the wall that becomes the outer curve. We therefore constructed five different protein variants of Bd1075 which were each C-terminally fused to mCherry (Fig. 6b). These constructs were then introduced into the *B. bacteriovorus* HD100 curved wild-type strain to generate single-crossover strains containing two copies of *bd1075*: the original wild-type and the new mCherry fusion. The sub-cellular localization of each fusion protein was then examined by epifluorescence microscopy.

As expected, the full-length protein localized to (what becomes) the convex cell face (Fig. 6c) and we noted no morphological nor deleterious effects resulting from the presence of two functional copies of *bd1075*. The LD-CPase domain contains three conserved catalytic triad residues: His-141, Ala-142 and Cys-156. Mutation of Cys-156 to alanine (C156A) in full length Bd1075 did not abrogate localization, indicating that LD-CPase activity is not involved in targeting (Fig. 6c).

As the Bd1075 C-terminal NTF2 domain (aa 196-304) is from a very broad protein superfamily^33^, it was not possible to predict a putative function for this domain but the structure with the P107/K108 loop bound over the NTF2 pocket (Fig. 5c) suggested that this was a substrate interaction mimic and a means to destabilize putative PG substrate interactions. We tested a role for the NTF2 domain in protein targeting via the generation of three additional mCherry fusions: (1) Full-length Bd1075 containing an NTF2 domain point mutation changing Tyr-274 (which forms the base of the substrate-binding pocket, Fig. 5c) to alanine: Y274A; (2) Bd1075 truncated protein comprising residues 1-E302 (similar to the shorter Csd6 and Pgp2) which terminates 2 residues prior to the completion of the NTF2 domain at A304, omitting the highly conserved W303 which the crystal structure had suggested to be a Bd1075-unique feature; and (3) Bd1075 truncated protein comprising residues 1-A304, completing the NTF2-like domain and including W303.

The Y274A and E302 truncated mutants (which both contain disruptions to the NTF2 domain) failed to localize to the outer curve, however the A304 truncation mutant which contains a complete NTF2 domain (and *Bdellovibrio*-specific residue W303) was correctly targeted (Fig. 6c). These results strongly supported the hypothesis that the NTF2 domain is involved in protein targeting to the PG side-wall (which becomes the convex cell face), and that the *Bdellovibrio*-specific NTF2 extension, including W303, is important for this function.

### The role of protein targeting in the generation of curvature

To investigate whether correct protein targeting is absolutely required to generate curvature, the five mCherry fusion constructs were introduced into the rod-shaped Δ*bd1075* mutant to generate single-crossover strains expressing solely the mCherry-tagged copy of *bd1075*. The sub-cellular localization and curvature of each fusion strain were examined. The full-length protein localized to (what becomes) the outer curve and completely complemented the curvature of Δ*bd1075* (Fig. 6d-e). The LD-CPase catalytic domain point mutant C156A did not restore curvature and remained localized at the center of the rod-shaped cell (Fig. 6d-e). Critically, neither of the NTF2 domain mutants Y274A nor E302 were correctly targeted and neither protein could complement the curvature of Δ*bd1075* (Fig 6d-e), highlighting the importance of the NTF2 domain and residue W303.

Together, these targeting data reveal that the NTF2 domain is responsible for targeting Bd1075 to the outer curve of the bacterial periplasm and that this specific localization is required to generate cell curvature.

## Discussion

In this work, we elucidate the first vibrioid cell shape-determinant of predatory *Bdellovibrio bacteriovorus* bacteria and show that vibrioid morphology confers dual fitness benefits to predators: rapid prey invasion and optimal intracellular growth. These findings contribute to fundamental knowledge of bacterial cell shape and deepen our understanding of the predatory process, which may assist the application of predatory bacteria as a therapeutic.

In vibrioid bacteria, intermediate filament-like (IF-like) cyto-or periskeletal elements frequently determine cell shape^24, 26^. IF-like proteins often contain coiled-coil rich protein (Ccrp) domains, however deletion of the sole *B. bacteriovorus* Ccrp protein was previously found not to affect vibrioid cell shape^34^. Here, we discover that the vibrioid curvature of *B. bacteriovorus* is instead determined by a PG cell wall hydrolase: Bd1075. Refining the initial prediction of an LDT domain, we show via sacculus studies that Bd1075 functions as an LD-CPase, cleaving both cross-linked and uncross-linked tetrapeptides to tripeptides in the predator PG cell wall (Fig. 4 and Table 1). This enzymatic activity was also observed for LD-CPase helical shape-determining proteins Csd6 (*H. pylori*) and Pgp2 (*C. jejuni*), highlighting the importance of biochemical validation of predicted LDT domains.

Considering how PG hydrolytic activity generates bacterial cell curvature, it is possible that targeted LD-CPase-mediated reduction in tetrapeptides may reduce localized cross-linking by DD- or LD-transpeptidases. Consistent with this idea, the overall peptide cross-linkage was slightly higher in cells lacking *bd1075* (64.4% compared to 61.1% in wild-type, Table 1). We theorize that a reduction in PG cross-linkage could soften one side of the PG sacculus such that the cell bulges slightly and becomes deformed by internal cellular turgor pressure which pushes outwards to generate an outer convex curve that may be fixed by subsequent, as yet uncharacterized, enzyme activity.

*B. bacteriovorus* vibrioid curvature is widely conserved within this group of invasive predators with the exception of the rod-shaped and long-cultured laboratory strain *B. bacteriovorus* 109J. Originally named *B. bacteriovorus* 109, the strain was re-designated as 109J following the observation that, in one research laboratory, predator cells had transitioned from a curved to non-vibrioid shape^35^. It is possible that long-term laboratory culture conditions in which prey are highly abundant may have removed the selection pressure for vibrioid morphology, resulting in the lab-evolved 57-residue deletion that we detect to have inactivated Bd1075_109J_. Bd1075_109J_ was not catalytically active upon PG (Supplementary Fig. 8a, Supplementary Fig. 9b, and Supplementary Table 1) and therefore could not complement curvature when cross-expressed in HD100 Δ*bd1075* (Supplementary Fig. 4). Despite retaining LD-CPase catalytic residues and targeting capability, the 57-residue N-terminal truncation could severely disrupt Bd1075_109J_ protein folding and therefore function. The mutation most likely occurred via homologous recombination between 8 bp repeats which flank the deleted region (Supplementary Fig. 3b); this has been observed in other predator genes^36^.

Bacterial morphology is evolutionarily conserved and known to confer selective advantages to different bacterial lifestyles^37, 38^. The helical morphology of *H. pylori*, for example, facilitates efficient bacterial motility through the gastric mucosa to allow pathogenic colonisation of the gastrointestinal tract^19^. Moreover, rod-shaped mutants of helical *C. jejuni* are deficient or have reduced fitness in a model of chick colonisation^21, 22^. In *V. cholerae*, vibrioid cell curvature (which is generated via a different non-enzymatic mechanism) increases motility through dense soft-agar matrices and promotes the colonisation of *V. cholerae* in motility-dependent mouse and rat infection models^26^.

Here, we propose new phenotypic roles for bacterial cell curvature: invasion and growth within Gram-negative prey bacteria. Straight rod-shaped *B. bacteriovorus* Δ*bd1075* predators invade prey significantly more slowly than curved wild-type predators (Fig. 2c). During prey invasion, *B. bacteriovorus* must presumably overcome opposing physical forces exerted upon itself by the turgid prey membrane and cell wall. Curved predators may distribute opposing forces as a glancing blow along the predator cell body, facilitating an efficient, curved trajectory into rounded prey, in contrast to force-intensive ‘head-on’ invasions by rod-shaped predators.

Unlike wild-type curved *B. bacteriovorus*, non-vibrioid Δ*bd1075* predators stretch and deform the rounded prey bdelloplast (Fig. 3c-e). Intriguingly, non-vibrioid Δ*bd1075* predators become gradually curved during elongation inside spherical prey bdelloplasts, despite the absence of *bd1075* (Fig. 3a-b). The Δ*bd1075* predator does not curve as tightly as the wild-type, however, and released progeny cells are not curved but rod-shaped, indicating that adoption of an intracellular curvature is temporary in this mutant (Fig. 3a). In wild-type curved predators, Bd1075 may thus be acting enzymatically on the *B. bacteriovorus* PG wall to tighten and “fix” a bdelloplast-imposed curve, consequently avoiding damage to the replicative niche, while also preparing curved and invasively-streamlined progeny predators for prey exit. This suggests a sensing and usage by predatory bacteria of the spherical prey environment - possibly to localize shape-determining enzyme(s) at a topologically-imposed curve (“curvature-templating”) and initiate the permanent “fixation” of a vibrioid cell shape. This topological-sensing could be a mechanism utilised more generally by bacteria or alternatively the structural differences of Bd1075 (monomeric, C-terminal extension) compared to the shape-determinants of free-living helical bacteria could render the usage of “curvature-templating” predator-specific.

Fluorescently-tagged Bd1075 specifically localizes to (what becomes) the outer convex face of vibrioid *B. bacteriovorus* cells (Fig. 6a). Most cell shape enzymes (for which localization is known) are found at the inner concave face of bacteria. These include the *C. crescentus* cytoskeletal polymer crescentin^24^ and *Vibrio cholera*e CrvA and CrvB which form periskeletal polymers^26, 39^. Only Bd1075 and the *H. pylori* bactofilin CcmA have been identified at the outer convex face of bacterial cells, occupying the periplasmic and cytoplasmic compartments, respectively^40^.

To our knowledge, there is currently no known mechanistic basis for asymmetric bacterial protein targeting to one lateral side-wall of the cell. Intrigued by this and guided by the phenotyping of Bd1075 protein truncations and point mutations, we discover that the extended C-terminal NTF2 domain (including the unique pocket residue W303) targets the protein to the convex face and is necessary to generate cell curvature (Fig. 6c-e). NTF2 is a nuclear envelope protein which transports molecules into eukaryotic nuclei^41^. The NTF2-like domain superfamily, however, comprises hundreds of thousands of proteins spanning the 3 domains of life and is associated with over 200 biological pathways, suggestive of divergent evolution^33, 42^. The NTF2 domain of *C. jejuni* Pgp2 was recently found by NMR studies to bind a variety of PG fragments, with specific secondary structure features shifting upon complexation^43^. The general agreement of some of the structure of monomeric Bd1075 with dimeric Pgp2 is suggestive that Bd1075 binds PG. The hydrogen-bond rich nature of the P107/K108 loop we observe in our crystal structure could be indicative of such an interaction by mimicking part of a PG muropeptide. Since the Pgp2 study invoked an induced fit on binding PG, it is particularly interesting that the third helix of the Bd1075 NTF2 domain (nH3, aa 233-248) and associated loop (labeled “lip”, aa 226-232) are shifted in relation to both Pgp2 and Csd6 (Fig. 5d). This helix was the major NTF2 feature that was reported to shift upon Pgp2-PG binding^43^. We postulate that its position has been modified in our Bd1075 structure by sidechains contacting the P107/K108 loop and may thus represent a “bound” state. There is also the potential for NTF2:LD-CPase domain crosstalk, given that a small LD-CPase domain helix (aa 125-132) shifts in response to NTF2 alterations^43^, and in Bd1075 appears to influence the disorder of the adjacent active site lid domain.

Since Bd1075 localizes to one lateral side-wall, one could theorize that the PG or outer membrane properties of that particular side-wall must be uniquely different to the other. One possibility is that the NTF2 domain recognizes a modification or substrate which is more abundant at this side-wall. Alternatively, the NTF2 domain may recognize the temporary physical curvature imposed by growth inside the spherical bdelloplast (“curvature-templating”) and direct Bd1075 to this curved cell face. Bd1075 may then exert LD-CPase activity, initiating the “fixation” of *B. bacteriovorus* curvature. Such a possible curvature-templating mechanism is supported by the localization of Bd1075 (C156A)-mCherry in Δ*bd1075* (Fig. 6d). This construct should be capable of correct asymmetric localization to one lateral side-wall (but not curvature generation), however the C156A-mCherry protein remained localized to the cytoplasm of the rod-shaped Δ*bd1075* attack-phase cells which have been released from bdelloplasts (Fig. 6d). This supports the idea that Bd1075 may sense and localize to temporary curvature that is templated by the spherical bdelloplast shape. These questions - which would present a significant investigative challenge beyond the scope of this study - could indicate whether bacteria have evolved to sense the topology of their physical space environment and use that dynamically to template their final morphotype during growth and replication.

Collectively, these findings advance our understanding of factors affecting the fitness of therapeutically-promising predatory bacteria and provide new mechanistic insights into the evolutionary importance of bacterial cell morphology.

## Methods

### Bacterial strains and culture

*Bdellovibrio bacteriovorus* strains were predatorily cultured in liquid Ca/HEPES buffer (5.94 g/l HEPES free acid, 0.284 g/l calcium chloride dihydrate, pH 7.6) and on solid YPSC overlays, both containing *Escherichia coli* S17-1 prey as described previously^44^. Where appropriate, kanamycin or gentamicin were added to growth media at concentrations of 50 µg ml^-1^ or 5 µg ml^-1^, respectively.

### Plasmid and strain construction

Primers and plasmids used in to construct strains used in this study are detailed in Supplementary Table 3 and Supplementary Table 4. Strains are listed in Supplementary Table 5. Bd1075 protein residue numbering was updated to correct for a previously mis-annotated start codon; the probable true start begins 2 codons downstream from that originally predicted (Supplementary Fig. 1b). To construct a markerless deletion of *bd1075*, 1 kb of upstream and downstream DNA were cloned into the suicide vector pK18*mobsacB* by Gibson assembly^45^ using the NEBuilder HiFi DNA Assembly Cloning Kit (New England Biolabs). The vector was introduced into *B. bacteriovorus* via an *E. coli* S17-1 conjugal donor strain and a double-crossover deletion mutant was isolated by sucrose suicide counter-selection as described previously^8, 46^. Deletion of *bd1075* was verified by RT-PCR, Sanger sequencing and whole-genome sequencing.

Strains for complementation tests were constructed by inserting the *bd1075* gene from either strain HD100 or strain 109J (gene ID: EP01_15440) plus the respective native gene promoter into the vector pMQBAD, a derivative of pMQ414 which expresses the fluorescent tdTomato protein and is capable of autonomous replication in *B. bacteriovorus*^47^. Constructs for complementation tests were verified by Sanger sequencing, conjugated into *B. bacteriovorus* and maintained under a gentamicin selection pressure.

To initially fluorescently-tag Bd1075, the stop codon of *bd1075* was replaced with the *mCherry* coding sequence to generate an in-frame C-terminal fusion terminating at the stop codon of the mCherry protein. The fluorescent strain was constructed and verified analogously to the Δ*bd1075* mutant to generate a markerless double-crossover Bd1075-mCherry strain. For more extensive analysis of domain functions, single-crossover mCherry fusions to either full length or truncated versions of Bd1075 were made. These were constructed by cloning 1 kb of upstream DNA, the DNA encoding each desired Bd1075 domain-test (minus the stop codon) and the *mCherry* sequence into pK18*mobsacB*. Point mutations in such constructs were generated using Q5^®^ Site-Directed Mutagenesis (New England Biolabs). Each construct was then conjugated into *B. bacteriovorus* to generate single-crossover merodiploid strains which were confirmed by Sanger sequencing and maintained under a kanamycin selection pressure.

### Protein expression and purification

The *bd1075* gene from *B. bacteriovorus* HD100, minus the signal peptide and stop codon, was cloned into the vector pET41 in-frame with a C-terminal histidine tag and transformed into *E. coli* BL21. *E. coli* BL21 was cultured in TB media and incubated at 37 °C with orbital shaking at 200 rpm to an OD_600_ of 0.6-0.8. *bd1075* expression was then induced with 0.5 mM IPTG at 18 °C for 16 h. Bd1075 was purified to near homogeneity using Ni-NTA affinity and size-exclusion chromatography, then dialyzed into either buffer containing 10 mM Na Citrate pH 6.0, 30 mM KCl, and 2% w/v glycerol (activity assays) or buffer containing 20 mM Na Citrate pH 6.0, 200 mM NaCl and 2 mM β-mercaptoethanol (structure studies).

### Muropeptide analysis

To culture *B. bacteriovorus* for sacculi isolation, 1 l of each *B. bacteriovorus* strain was grown on either *E. coli* S17-1 or *E. coli* S17-1 pUC19 (gentamicin-resistant prey for complementation test strains containing the pMQBAD plasmid). Following complete predatory lysis of *E. coli* prey during a 24 h incubation at 29 °C with orbital shaking at 200 rpm, *B. bacteriovorus* attack-phase cultures were passed through a 0.45 µm filter to remove any remaining prey debris. To culture *E. coli* BW25113 for sacculi isolation, *E. coli* was grown at 37 °C for 16 h with orbital shaking at 200 rpm. Cultured *B. bacteriovorus* or *E. coli* cells were then centrifuged at 15,000 g for 30 min at 4 °C, resuspended in 6 ml of ice-cold PBS, and then boiled in 6 ml of 8% SDS for 30 min to lyse the cells and liberate sacculi. Peptidoglycan was purified from the cell lysates, then muropeptides in the supernatant were reduced with sodium borohydride and HPLC analysis performed as previously described^48^.

To test the *in vitro* activity of Bd1075_HD100_, sacculi from either *B. bacteriovorus* HD100 Δ*bd1075*, *B. bacteriovorus* wild-type 109J or *E. coli* BW25113 were incubated with 10 µM of *B. bacteriovorus* Bd1075_HD100_ in 50 mM Tris-HCl, 50 mM NaCl, pH 7.0 for 16 h at 37 °C on a thermomixer at 900 rpm. The control sample received no enzyme. To stop Bd1075 activity, the samples were boiled at 100 °C for 10 min and an equal volume of 80 mM sodium phosphate, pH 4.8, was added. The samples were incubated with 10 µg of cellosyl (Hoechst, Frankfurt am Main, Germany) for a further 16 h at 37 °C on a thermomixer at 900 rpm, boiled for 10 min and centrifuged at room temperature for 15 min at 16,000 g. Muropeptides present in the supernatant were reduced with sodium borohydride and analyzed by HPLC as described^48^.

### Structure determination

Purified Bd1075 at ∼25 mg/ml was used for screening. Crystals were grown at 18 °C using the sitting drop technique in 4 μl drops composed of equal volumes of protein and reservoir solution. B1075 crystals were obtained in the BCS™ screen (Molecular Dimensions) in condition 2-44, comprising 0.1 M Tris pH 7.8, 0.1 M KSCN, 0.1 M NaBr and 25% PEG Smear broad range. Crystals were cryo-protected in mother liquor supplemented with 25% (v/v) ethylene glycol and flash cooled in liquid nitrogen. Diffraction data were collected at the Diamond Light Source in Oxford, UK (Supplementary Table 2). Data reduction and processing were completed using XDS^49^ and the xia2 suite^50^. Bd1075 phasing was achieved using a merged SAD data set (9000 frames, 0.1° oscillations) collected at a wavelength of 0.91 Å corresponding to the bromide anomalous scattering peak. The collected data were input into CCP4 online CRANK2^51^, which located six bromide sites with an initial FOM of 0.14, followed by iterative cycles of building and model-based phasing improvement. The obtained model was further built and modified using COOT^52^, with cycles of refinement in PHENIX^53^.

### Phase-contrast and epifluorescence microscopy

*B. bacteriovorus* cells were immobilized on a thin 1% Ca/HEPES buffer agarose pad and visualized under a Nikon Ti-E inverted epifluorescence microscope equipped with a Plan Apo 100x Ph3 oil objective lens (NA: 1.45), an mCherry filter (excitation: 555 nm, emission: 610-665 nm), and an mCerulean3 filter (excitation: 440 nm, emission: 470-490 nm). Images were acquired on an Andor Neo sCMOS camera with Nikon NIS software. Images were analyzed using the Fiji distribution of ImageJ^54^ and minimally processed using the sharpen and smooth tools, with adjustments to brightness and contrast. The MicrobeJ plug-in for ImageJ^55^ was used to measure cell morphologies and detect fluorescent foci. *B. bacteriovorus* attack-phase cells were generally identified by the parameters of area: 0.2-1.5 µm^2^, length: 0.5-2.5 µm, width: 0.2-0.8 µm, and circularity: 0-0.9 A.U. For the detection of Bd1075-mCherry fluorescent fusions in a curved wild-type genetic background, *B. bacteriovorus* cells were defined with the same parameters, but excluded cells with a curvature of <0.6 so as to only measure localization in cells with a definitively curved shape. Curvature parameters were set to 0-max to allow measurements of curvature for fluorescent fusions expressed in a non-curved Δ*bd1075* genetic background. Fluorescent Bd1075-mCherry foci within attack-phase *B. bacteriovorus* cells were detected with the foci method using default maxima settings and an association with parent bacteria with a tolerance of 0.1 µm. Images were manually inspected to ensure cells had been correctly detected before measurements were acquired. To analyze prey bdelloplast morphology, bdelloplasts were generally identified by the parameters of area: 1.0-max µm^2^, length: 0.5-max µm, width: 0.5-max µm, and circularity: 0.8-max A.U. *B. bacteriovorus* cells were detected in the maxima channel with the bacteria method using default maxima settings and an association with parent bacteria with a tolerance of 0.1 µm. Only the morphologies of prey bdelloplasts which contained a single *B. bacteriovorus* predator were measured.

### Electron microscopy

*B. bacteriovorus* cells were cultured for 24 h, then concentrated by microcentrifugation at 5,000 g for 10 min followed by careful resuspension in 1 ml of Ca/HEPES. *B. bacteriovorus* cells were applied to glow-discharged Formvar/Carbon-coated 200-mesh copper grids (EM Resolutions), stained with 0.5% uranyl acetate for 1 min, then de-stained with Tris-EDTA pH 7.6 for 30 s. Samples were imaged under a FEI Tecnai G2 12 Biotwin transmission electron microscope at 100 kV.

### Time-lapse microscopy

For time-lapse microscopy, 1 ml cultures of attack-phase *B. bacteriovorus* and 50 µl of stationary-phase *E. coli* S17-1 were microcentrifuged separately at 17,000 g for 2 min, then resuspended in 50 µl of Ca/HEPES. Predators and prey were then mixed together and immediately transferred to a thin 0.3% Ca/HEPES agarose pad. Cells were visualized under a Nikon Eclipse E600 upright microscope equipped with a 100x oil objective lens (NA: 1.25) and a Prior Scientific H101A XYZ stage which allowed 6 specific fields of view to be revisited over the time-lapse sequence. Image frames were captured every 1 min for at least 2 h on a Hammamatsu Orca ER Camera with Simple PCI software. Time-lapse videos of *B. bacteriovorus* attaching to and entering *E. coli* prey were analyzed in Simple PCI software. Attachment time was measured by counting the number of frames (1 frame = 1 min) between initial irreversible predator attachment to prey and the first indication of the predator moving into prey. Entry time was measured by counting the number of frames between the first indication of the predator moving into prey and the predator residing completely inside the bdelloplast.

### Statistical analysis

Statistical analysis was performed in Prism 8.0 (GraphPad). Data were first tested for normality and then analyzed using an appropriate statistical test. The number of biological repeats carried out, n values for cell numbers, and the statistical test applied to the data set are described in each figure legend.

### Data availability

Coordinates and structure factors have been deposited in the PDB under the accession code 7O21.

## Acknowledgements

This work was funded by a Wellcome Trust PhD studentship (215025/Z/18/Z) to E.J.B., a Becas Chile studentship (72180329) to M.V-D., a BBSRC studentship to A.W. and the UKRI Strategic Priorities Fund (https://www.ukri.org) EP/T002778/1 to W.V. R.E.S, C.L and A.L.L are currently funded by a Wellcome Trust Investigator Award in Science (209437/Z/17/Z). We thank Chloe Hudson for trials of Bd1075 purification, Daniela Vollmer for purification of PG, Rob Till for general laboratory support, and Block Allocation Group mx19880 for access to Diamond Synchrotron. Genome sequencing was provided by MicrobesNG (http://www.microbesng.uk) which is supported by the BBSRC (grant number BB/L024209/1). Electron microscopy was carried out at the Nanoscale and Microscale Research Centre at the University of Nottingham.

## Author contributions

E.J.B. made all deletion, complementation and fluorescent strains of Bd1075, carried out curvature analysis and conducted electron, time-lapse, and epifluorescence microscopy experiments with supervision by R.E.S. and C.L. J.B. carried out PG sacculi purifications and HPLC analysis with supervision by W.V. I.T.C. made the *bd1075* expression construct for protein purification. A.W purified the protein with purification optimization and protocol design from I.T.C. M.V-D crystallized and solved the structure of Bd1075. All protein work was supervised by A.L.L. E.J.B., R.E.S. and A.L.L. wrote the manuscript and all co-authors read and approved the final manuscript.

## Competing interests

The authors declare no competing interests.

## Additional information

Supplementary information is available at [X]

Correspondence and requests for materials should be addressed to R.E.S.

## Notes

### Competing Interest Statement

The authors have declared no competing interest.

