## Supplementary information for "Asymmetric peptidoglycan editing generates the curvature of predatory bacteria, optimizing invasion and replication within a spherical prey niche"

Banks *et al*.

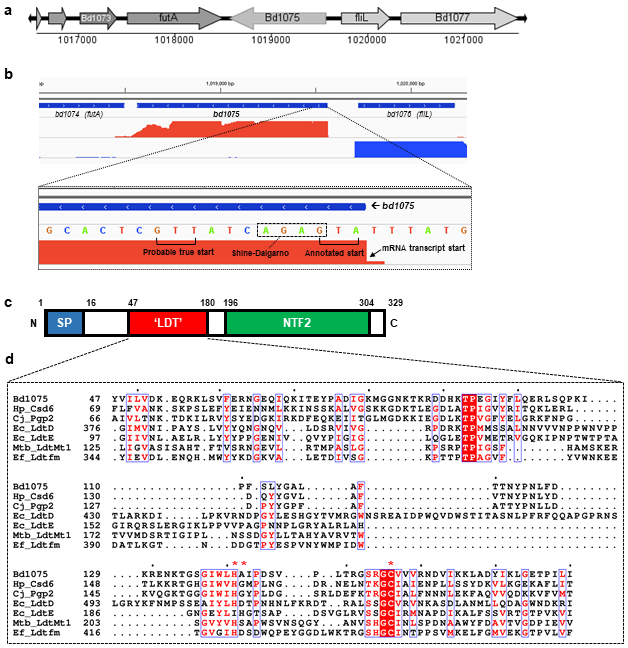

**Supplementary Fig. 1 Bioinformatic predictions of Bd1075.**

**a** Genomic location of *bd1075* in the *B. bacteriovorus* HD100 genome, viewed in xBASE^1^. The *bd1075* gene is monocistronic and flanked by divergently transcribed genes. **b** RNA-Seq reads^2^ aligned to the *B. bacteriovorus* HD100 genome with Rockhopper^3, 4^ and visualized in Integrated Genomics Viewer^5^. The DNA sequence has been reverse-complemented for ease of viewing. The mRNA transcript of *bd1075* begins at the start codon, indicating that the start codon of the gene was probably mis-annotated. The probable Shine-Dalgarno sequence (GAGA) and true start codon (TTG) are annotated. Correcting the start site annotation removes the first 3 residues (MRL) of Bd1075 but does not affect any signal peptide or domain predictions. **c** Schematic of the predicted domain structure of Bd1075. Numbers indicate residue position; SP: signal peptide; ‘LDT’: predicted LD-transpeptidase-family domain; NTF2: nuclear transport factor 2-like domain. **d** Multiple sequence alignment of the *B. bacteriovorus* HD100 Bd1075 LDT domain against the LDT domains (in descending order) of: *Helicobacter pylori* Csd6, *Campylobacter jejuni* Pgp2, *Escherichia coli* LdtD, *Escherichia coli* LdtE, *Mycobacterium tuberculosis* Ldt_Mt1_, and *Enterococcus faecium* Ldt_fm_. Catalytic triad residues are indicated by red asterisks. The alignment was generated in Clustal Omega^6^ and visualized with ESPript 3^7^.

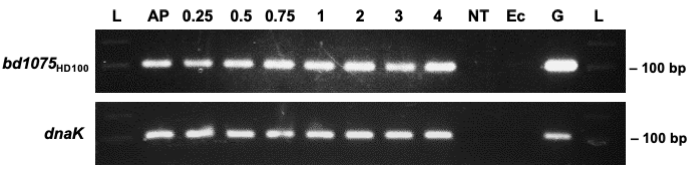

**Supplementary Fig. 2 *bd1075*_HD100_ is constitutively expressed during the predatory cycle.**

Reverse-transcriptase PCR was performed on *B. bacteriovorus* HD100 RNA isolated at timepoints throughout the predatory cycle using primers designed to amplify a 102 bp product internal to the *bd1075* gene. L: 100 bp NEB molecular weight ladder; AP: attack-phase; 0.25-4: hours since the start of predation; NT: no template RNAase-free water control, Ec: *E. coli* S17-1 RNA control, G: *B. bacteriovorus* HD100 genomic DNA positive control. Both *bd1075* and the control gene *dnaK* are constitutively expressed throughout predation. Two independent biological repeats were carried out.

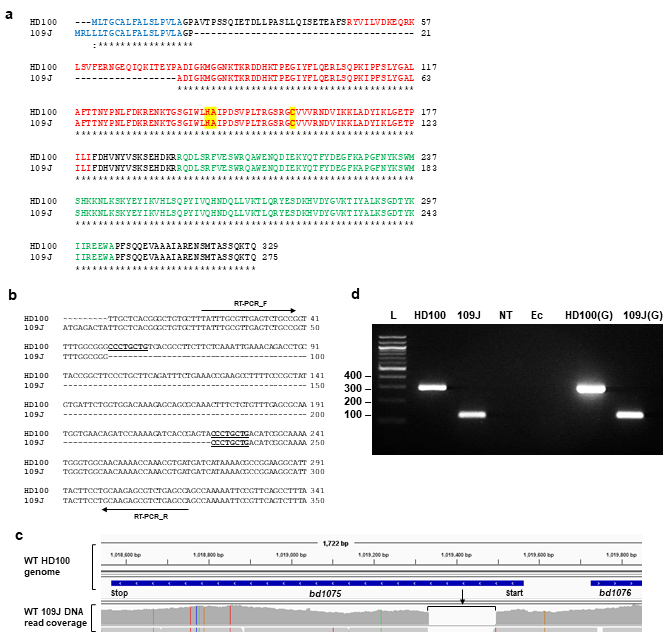

**Supplementary Fig. 3 *bd1075* is expressed in strain 109J and contains an N-terminal 57 residue truncation.**

**a** Pairwise amino acid alignment of wild-type Bd1075_HD100_ and wild-type Bd1075_109J_. Asterisks indicate identical protein residues and dashes indicate the 57 amino acid truncation present in Bd1075_109J_ between Proline-18 and Tyrosine-74. Blue residues: signal peptide; red residues: predicted ‘LD-transpeptidase (LDT) domain; red residues highlighted in yellow: LDT catalytic triad residues; green residues: NTF2 domain. **b** Pairwise DNA alignment of wild-type HD100 *bd1075*_HD100_ and wild-type *bd1075*_109J_ from 1-350 bp, showing the truncation (indicated by dashes) present in strain 109J which is flanked by 8 bp flanking repeats (underlined and emboldened). **c** Wild-type 109J DNA sequence reads mapped to the wild-type HD100 genome, showing the truncation present in *bd1075*_109J_. Data were visualized in Integrated Genomes Viewer. **d** Reverse-transcriptase PCR was performed on RNA isolated from attack-phase wild-type HD100 and wild-type 109J using primers (indicated with black arrows in (**b**) designed to anneal to either side of the *bd1075*_109J_ truncation. Expected product sizes of 298 bp (HD100) and 127 bp (109J) confirmed the presence of the 109J truncation in the RNA transcript. L: 100 bp NEB molecular weight ladder; HD100 & 109J: RNA isolated from strains HD100 and 109J; NT: no template RNAase-free water control; Ec: *E. coli* S17-1 RNA control; HD100(G) and 109J(G): *B. bacteriovorus* genomic DNA positive controls. Two independent biological repeats were carried out.

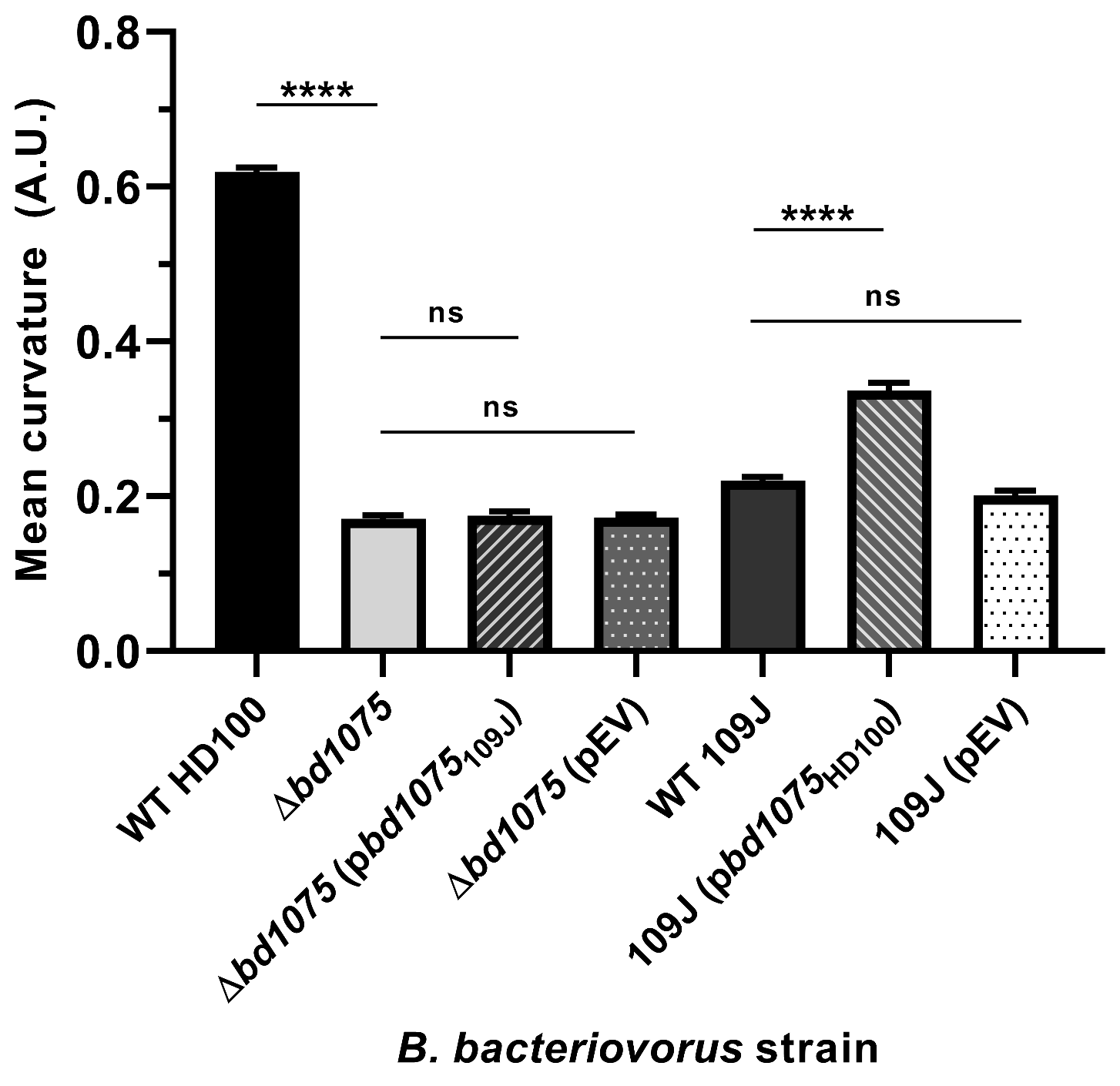

**Supplementary Fig. 4 Curvature of *B. bacteriovorus* 109J and additional complementation strains.**

Curvature measurements of *B. bacteriovorus* attack-phase cells. n = 759-2503 cells per strain from 3 biological repeats. WT HD100, Δ*bd1075* and Δ*bd1075* (pEV) data are reproduced from Fig. 1c. Error bars represent standard error of the mean. ns: non-significant (p>0.05), ****: p<0.0001; Kruskal-Wallis test. Frequency distributions are included in Supplementary Fig. 5b.

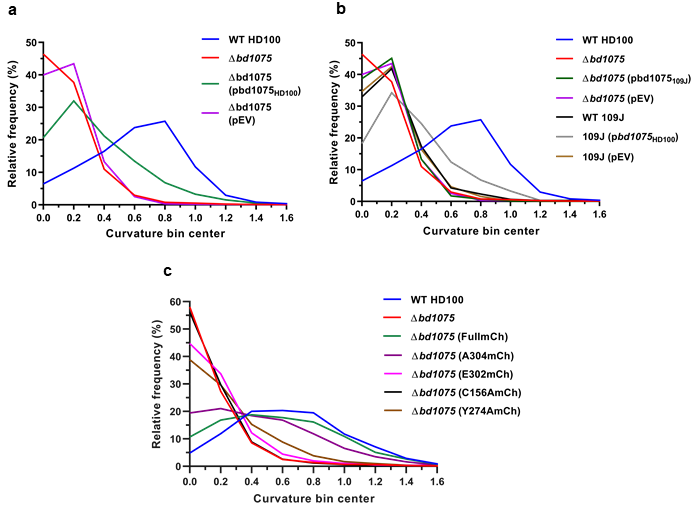

**Supplementary Fig. 5 Frequency distributions of *B. bacteriovorus* cell curvature.**

Curvature frequency distributions of attack-phase *B. bacteriovorus* strains shown in Fig. 1c (**a**), Supplementary Fig. 4 (**b**) and Fig. 6e (**c**). Graphs show the relative percentage of cells that have a particular value of curvature. n = 1920-2503 (**a**), n = 759-2502 (**b**) and n = 1886-2812 (**c**) total cells per strain from 3 biological repeats. Bin width = 0.2.

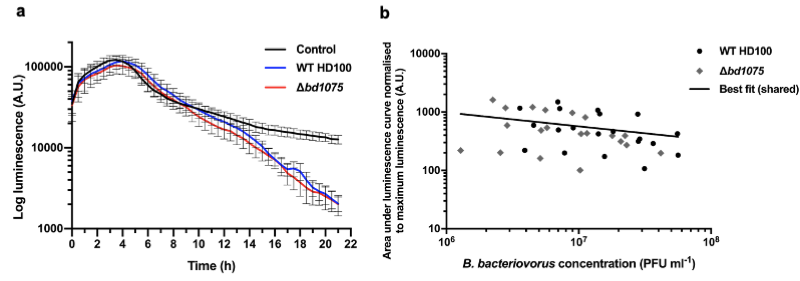

**Supplementary Fig. 6 Predation on *E. coli* in liquid culture by *B. bacteriovorus* HD100 wild-type and Δ*bd1075*.**

Comparison of the predation efficiency of *B. bacteriovorus* WT HD100 and Δ*bd1075* in liquid culture. Predation efficiency was quantified by the rate in the reduction of *E. coli* S17-1 prey luminescence, measured every 30 min for 21 h. **a** Representative luminescence prey-death curve. Blue line: wild-type HD100; red line: Δ*bd1075*; black line: control of heat-killed predator cells. Error bars represent standard error of the mean. **b** The area under each luminescence curve was measured and normalised to the maximum luminescence. These values were plotted against corresponding *B. bacteriovorus* predator concentrations which were enumerated by plaque counts. Black circles: wild-type HD100; grey diamonds: Δ*bd1075*. Both data sets could be analyzed by a shared non-linear regression line of best fit, indicating that there was no significant difference between strains (p=0.70). Data are from 5 biological repeats.

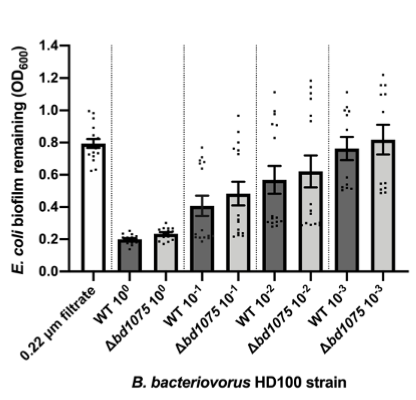

**Supplementary Fig. 7 Predation on *E. coli* biofilms by *B. bacteriovorus* HD100 wild-type and Δ*bd1075.***

Comparison of *B. bacteriovorus* WT HD100 and Δ*bd1075* predation upon pre-formed *E. coli* S17-1 biofilms within 96-well PVC microtiter plates. Predation efficiency was assessed by quantification of remaining *E. coli* biofilm (OD_600_) after 24 h incubation with each predator at a range of dilutions: neat (10^0^), 10^-1^, 10^-2^, and 10^-3^. 0.22 μm filtrate: *B. bacteriovorus* filtered through a 0.22 μm membrane to give a no-predator control. Data points represent 15 technical repeats from 3 independent biological repeats and error bars represent the standard error of the mean. There was a small significant difference (p=0.01; unpaired t-test) between neat WT and Δ*bd1075*, however the difference was not considered biologically meaningful as it was within the error margins of the experiment; predator plaque enumerations showed that the concentration of viable Δ*bd1075* that had been added was slightly lower than the WT. No other comparisons were significant (p>0.05; Mann-Whitney test).

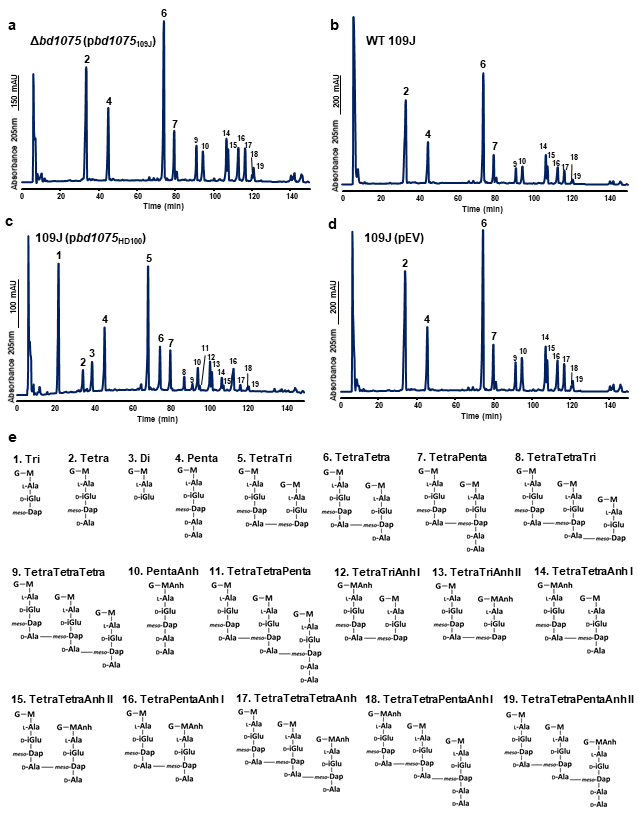

**Supplementary Fig. 8 Muropeptide analysis of *B. bacteriovorus* HD100 and 109J strains.**

**a-d** HPLC elution profiles of muropeptides released from peptidoglycan sacculi from attack-phase *B. bacteriovorus* cells of **a** HD100 ∆*bd1075* (p*bd1075*_109J_) - *bd1075*_109J_ expressed in ∆*bd1075* , **b** wild-type (WT) 109J, **c** 109J (p*bd1075*_HD100_) - *bd1075*_HD100_ expressed in 109J, and **d** 109J (pEV) - empty vector (EV) control in 109J. Representative chromatograms of 2 biological repeats are shown. **e** Proposed structures of muropeptides. Numbers correspond to those above peaks in **a-d**. G: *N*-acetylglucosamine, M: *N*-acetylmuramitol, MAnH: 1,6-anhydro-*N*-acetylmuramic acid, L-Ala: L-alanine, D-Glu: D-glutamic acid, *meso*-Dap: *meso*-diaminopimelic acid, D-Ala: D-alanine.

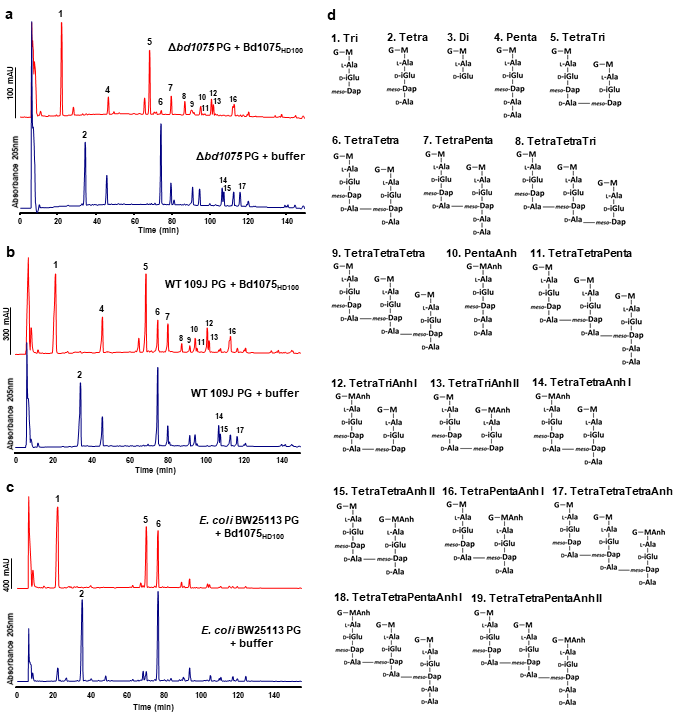

**Supplementary Fig. 9 Muropeptide analysis of peptidoglycan sacculi treated with purified Bd1075 *in vitro*.**

HPLC elution profiles of muropeptides released from peptidoglycan sacculi from **a** *B. bacteriovorus* ∆*bd1075*, **b** *B. bacteriovorus* wild-type 109J, and **c** *E. coli* wild-type BW25113 that had been treated with either purified Bd1075_HD100_ enzyme (top) or buffer control (bottom). Data are from 1 biological repeat. **d** Proposed structures of muropeptides. Numbers correspond to those above peaks in **a-c**. G: *N*-acetylglucosamine, M: *N*-acetylmuramitol, MAnh: 1,6-anhydro-*N*-acetylmuramic acid, L-Ala: L-alanine, D-Glu: D-glutamic acid, *meso*-Dap: *meso*-diaminopimelic acid, D-Ala: D-alanine.

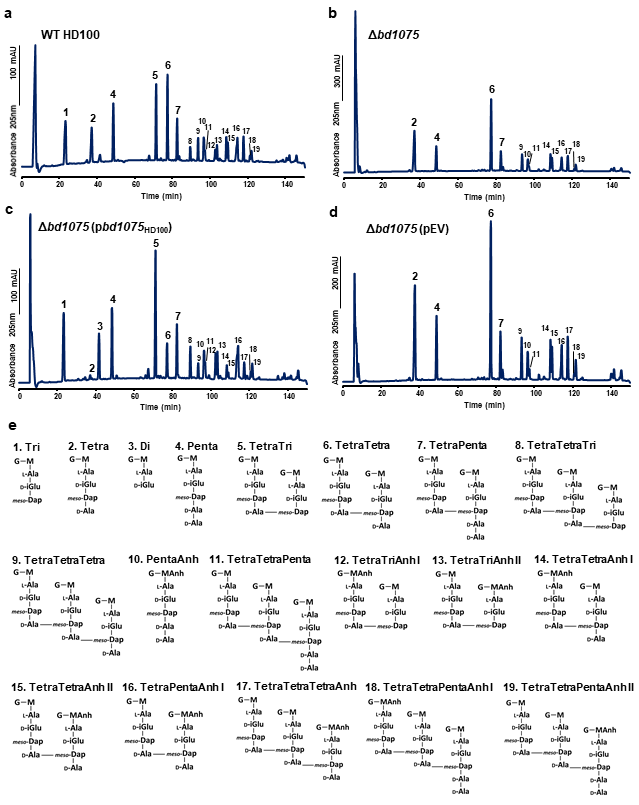

**Supplementary Fig. 10 Muropeptide analysis of *B. bacteriovorus* HD100.**

**a-d** HPLC elution profiles of muropeptides released from peptidoglycan sacculi isolated from attack-phase *B. bacteriovorus* cells of **a** wild-type (WT) HD100, **b** ∆*bd1075*, **c** ∆*bd1075* (p*bd1075*_HD100_) - *bd1075*_HD100_ expressed in ∆*bd1075*, and **d** ∆*bd1075* (pEV) - empty vector control in ∆*bd1075*. Representative chromatograms of 2 biological repeats are shown. **e** Proposed structures of muropeptides. Numbers correspond to those above peaks in **a-d**. G: *N*-acetylglucosamine, M: N-acetylmuramitol, MAnh: 1,6-anhydro-*N*-acetylmuramic acid, L-Ala: L-alanine, D-Glu: D-glutamic acid, *meso*-Dap: *meso*-diaminopimelic acid, D-Ala: D-alanine.

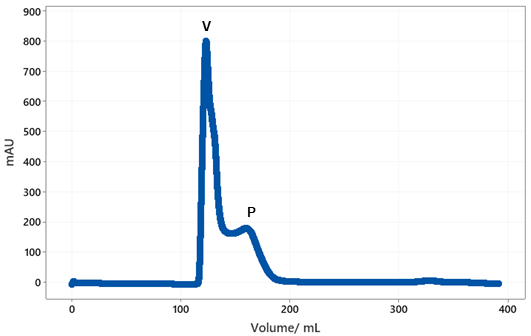

**Supplementary Fig. 11 Gel filtration profile of Bd1075 protein.**

Bd1075 distributes between two peaks on a Superdex 75 26/60 column. ‘V’ corresponds to the void volume of the column (elution volume of 120 ml) and ‘P’ to the elution volume (170 ml) of monomeric Bd1075. The monomeric peak of Bd1075 was pooled for crystallization and structure determination.

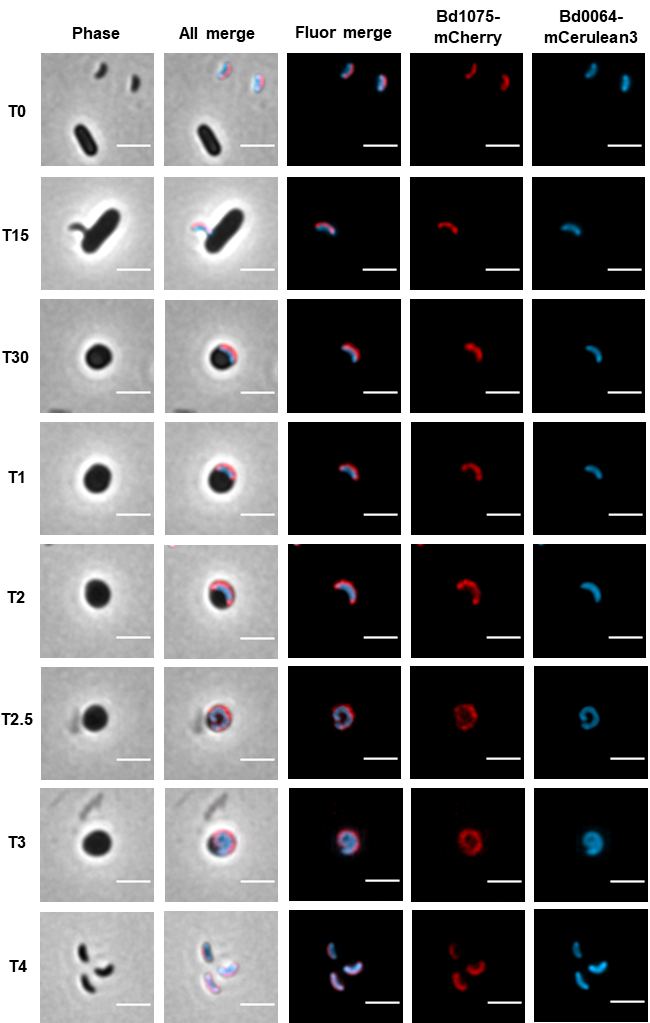

**Supplementary Fig. 12 Localization of Bd1075-mCherry during the predatory cycle of *B. bacteriovorus***

Growth of *B. bacteriovorus* HD100 containing chromosomal fusions of both Bd0064-Cerulean3 (to label the predator cytoplasm) and Bd1075-mCherry inside *E. coli* S17-1 pZMR100 prey bdelloplasts. T = min (0-30) or hours (1-4) elapsed since predators and prey were initially mixed. Scale bars = 2 µm. Images are representative of cells from 3 biological repeats.

**Supplementary Table 1. Quantification of muropeptides released from *B. bacteriovorus* HD100 and 109J sacculi.**

Values represent the relative percentage area of each muropeptide peak in **Supplementary Fig. 8**. For the strain HD100 ∆*bd1075* (p*bd1075*_109J_), numbers with an asterisk differ from WT HD100 values (Table 1) by more than 30% and numbers that are additionally emboldened differ by more than 50%. Values of strains 109J (p*bd1075*_HD100_) and 109J (pEV) are compared to WT 109J in identical manner.

| **Muropeptide** | **Relative peak area (%)^1^ in *B. bacteriovorus* strain** | | | |
| --- | --- | --- | --- | --- |
|  | **Δ*bd1075* (p*bd1075*_109J_)** | **WT 109J** | **109J (p*bd1075*_HD100_)** | **109J (pEV)** |
| **Monomers** |  |  |  |  |
| Tri | **n.d.*^2^** | n.d. | **18.7 ± 3.0*** | 0.3 ± 0.4 |
| Tetra | **22.2 ± 1.5*** | 28.0 ± 4.3 | **3.0 ± 0.8*** | 26.4 ± 0.5 |
| Di | **n.d.*** | n.d. | **6.2 ± 0.8*** | n.d. |
| Penta | 12.8 ± 5.9 | 13.1 ± 4.1 | 14.0 ± 1.9 | 14.6 ± 2.5 |
| Monomer anhydroMurNAc | 2.8 ± 4.0* | 2.5 ± 3.5 | 2.2 ± 3.2 | 3.1 ± 4.3 |
| Monomers (Total) | 35.1 ± 7.4 | 41.0 ± 8.4 | 42.0 ± 4.8 | 41.3 ± 2.7 |
| **Dimers** |  |  |  |  |
| TetraTri | **0.5 ± 0.7*** | 0.3 ± 0.4 | **26.2 ± 0.6*** | 0.2 ± 0.2* |
| TetraTetra | **33.4 ± 0.9*** | 33.8 ± 1.1 | **8.8 ± 0.8*** | 34.3 ± 1.5 |
| TetraPenta | 12.6 ± 0.5 | 11.4 ± 1.4 | 13.2 ± 0.5 | 12.1 ± 0.9 |
| Dimer anhydroMurNAc | 17.2 ± 1.3 | 15.1 ± 1.3 | 16.2 ± 1.1 | 15.2 ± 0.4 |
| Dimers (total) | 46.5 ± 0.6 | 45.5 ± 0.7 | 48.2 ± 1.0 | 46.6 ± 0.3 |
| **Trimers** |  |  |  |  |
| TetraTetraTri | **n.d.*** | n.d. | **2.3 ± 0.6*** | n.d. |
| TetraTetraTetra | **11.6 ± 2.6*** | 7.9 ± 1.9 | **2.5 ± 0.4*** | 7.2 ± 0.9 |
| TetraTetraPenta | 6.9 ± 5.4 | 5.6 ± 5.8 | 5.1 ± 4.8 | 4.9 ± 4.0 |
| Trimer anhydroMurNAc | 10.4 ± 2.8* | 6.6 ± 2.5 | **2.9 ± 1.0*** | 5.9 ± 0.7 |
| Trimers (total) | 18.4 ± 8.0 | 13.5 ± 7.7 | 9.8 ± 5.8 | 12.1 ± 3.0 |
| Dipeptides (total) | **n.d.*** | n.d. | **6.2 ± 0.8*** | n.d. |
| Tripeptides (total) | **0.2 ± 0.3*** | 0.1 ± 0.2 | **32.6 ± 3.1*** | **0.3 ± 0.5*** |
| Tetrapeptides (total) | 78.3 ± 4.0* | 79.3 ± 1.3 | **38.9 ± 2.3*** | 77.4 ± 0.3 |
| Pentapeptides (total) | 18.6 ± 0.3 | 18.1 ± 2.0 | 20.0 ± 3.2 | 19.2 ± 3.6 |
| AnhydroMurNAc (total) | 14.9 ± 2.4 | 12.3 ± 2.1 | 11.3 ± 3.4 | 12.6 ± 4.8 |
| Average chain length | 6.8 ± 1.1 | 8.3 ± 1.4 | 9.2 ± 2.8 | 8.5 ± 3.2 |
| Degree of cross-linkage | 35.5 ± 5.0 | 31.7 ± 5.5 | 30.7 ± 3.4 | 31.4 ± 1.9 |
| % peptides in cross-links | 64.9 ± 7.4 | 59.0 ± 8.4 | 58.0 ± 4.8 | 58.7 ± 2.7 |

^1^ values are mean ± variation of two biological replicates.

^2^ n.d., not detected.

**Supplementary Table 2. Bd1075 structure data collection and refinement statistics.**

|  | **Bd1075-Br SAD** | **Bd1075 native** |
| --- | --- | --- |
| **Data Collection** |  |  |
| Resolution range (Å) | 40.19 - 1.39 | 43.92 - 1.34 |
| Space group | P 2_1_ | P 2_1_ |
| Cell dimensions |  |  |
| *a, b, c (Å)* | 49.11, 100.39, 62.28 | 49.189, 100.631, 62.2787 |
| *α, β, γ (°)* | 90, 108, 90 | 90, 108.07, 90 |
| Total reflections | 593918 (14215) | 847431 (69189) |
| Unique reflections | 94881 (4170) | 123276 (11323) |
| Completeness (%) | 82.47 (36.26) | 95.37 (85.34) |
| Redundancy | 6.3 (3.4) | 6.9 (6.1) |
| <I>/σI | 20.58 (0.72) | 17.51 (0.49) |
| R_meas_ | 0.04208 (1.62) | 0.06161 (2.08) |
| R_merge_ | 0.03875 (1.39) | 0.05678 (1.9) |
| CC_1/2_ | 1.00 (0.36) | 0.999 (0.33) |
| CC_anom_ | 0.3 (0.0) |  |
| **Refinement** |  |  |
| Maximum resolution (Å) |  | 1.34 |
| Number of reflections used |  | 122860 (10948) |
| Number of reflections for R_free_ |  | 6092 (535) |
| R_work_ |  | 18.6 |
| R_free_ |  | 21.2 |
| RMSD |  |  |
| Bonds (Å) |  | 0.011 |
| Angles (°) |  | 1.02 |
| Number of non-hydrogen atoms |  |  |
| Protein |  | 4686 |
| Ligand/ion |  | 61 |
| Water |  | 504 |
| *B*-factors |  |  |
| Protein |  | 24.22 |
| Ligand/ion |  | 41.63 |
| Water |  | 34.37 |
| Ramachandran favoured (%) |  | 97.57 |
| Ramachandran allowed (%) |  | 2.43 |
| Ramachandran outliers (%) |  | 0.00 |
| Rotamer outliers (%) |  | 0.39 |

| **Supplementary Table 3. Primers used in this research study.** | | |
| --- | --- | --- |
| **Primer** | **Sequence (5’ 🡪 3’)** | **Purpose** |
| 1075_up_F | CGTTGTAAAACGACGGCCAGTGCCAGACCATCGGAACCGCGTG | Deletion of *bd1075* |
| 1075_up_R | CTAGTTTATTGCGTCTCATAAATACTATTATGCCCGAATAGGAC |  |
| 1075_down_F | AGTATTTATGAGACGCAATAAACTAGGCTGTAAAG |  |
| 1075_down_R | GGAAACAGCTATGACCATGATTACGAGCCAAGTTGGTTTTGTATTC |  |
| 1075_HD100c_F | cagctggtaccatatgggaattcgaACTTTATTTACATTTAAATTACACGG | Cloning *bd1075*_HD100_ gene for complementation |
| 1075_HD100c_R | cttctctcatccgccaaaacagccaTATCGAGTTGATGAAAAAAGC |  |
| 1075_109Jc_F | cagctggtaccatatgggaattcgaACTTTATTTACATTTAAATTACACGG | Cloning *bd1075*_109J_ gene for complementation |
| 1075_109Jc_R | cttctctcatccgccaaaacagccaGTATCGAGTTGATGAAAAAAG |  |
| Bd0064_F | cgttgtaaaacgacggccagtgccaGTGGAGGACACATATACAGTTC | Bd0064-mCerulean3 single-crossover fusion |
| Bd0064_R | cttgctcaccatTCCGACTTTTTTAAAGATCGTG |  |
| mCeru_F | taaaaaagtcggaATGGTGAGCAAGGGCGAG |  |
| mCeru_R | ggaaacagctatgaccatgattacgTTACTTGTACAGCTCGTCCATG |  |
| 1075_up_F | CGTTGTAAAACGACGGCCAGTGCCAGACCATCGGAACCGCGTG | Bd1075-mCherry full-length double-crossover fusion |
| 1075_mCh_up_R | CCTTGCTCACCATTTGCGTTTTCTGGGAAGAGG |  |
| 1075_mCh_F | CCAGAAAACGCAAATGGTGAGCAAGGGCGAG |  |
| 1075_mCh_R | TTTACAGCCTAGTTTACTTGTACAGCTCGTCCATG |  |
| 1075_mCh_down_F | GCTGTACAAGTAAACTAGGCTGTAAAGGCAAAAAAAAAG |  |
| 1075_down_R | GGAAACAGCTATGACCATGATTACGAGCCAAGTTGGTTTTGTATTC |  |
| 1075_up_genF | CGTTGTAAAACGACGGCCAGTGCCAGACCATCGGAACCGCGTG | Bd1075-mCherry truncation and point mutant fusions (single-crossover) |
| 1075_mCh_SXO_genR | GGAAACAGCTATGACCATGATTACGTTACTTGTACAGCTCGTCCATG |  |
| Full_1075_R | CTTGCTCACCATTTGCGTTTTCTGGGAAGAGG |  |
| Full_mCh_F | CCAGAAAACGCAAATGGTGAGCAAGGGCGAG |  |
| E302_1075_R | CTTGCTCACCATTTCTTCTCGGATGATTTTATAAGTGTCAC |  |
| E302_mCh_F | CATCCGAGAAGAAATGGTGAGCAAGGGCGAG |  |
| A304_1075_R | CTTGCTCACCATAGCCCATTCTTCTCGGATG |  |
| A304_mCh_F | AGAAGAATGGGCTATGGTGAGCAAGGGCGAG |  |
| C156A_F | CTCTCGCGGCGCGGTCGTTGTTCGTAACGACG |  |
| C156A_R | CCGCGGGTCAAAGGCACC |  |
| Y274A_F | CCTGCAGCGCGCGGAATCTGACAAAC |  |
| Y274A_R | GTTTTCACCAGCAACTGATC |  |
| RT_1075_F | CAACGATCAGTTGCTGGTG | RT-PCR primers for expression across predatory lifecycle |
| RT_1075_R | AAGTGTCACCGGACTTGAGG |  |
| RT_dnaK_F | TGAGGACGAGATCAAACGTG |  |
| RT_dnaK_R | AAACCAGGTTGTCGAGGTTG |  |
| RT_1075_GAP_F | TATTTGCGTTGAGTCTGCCG | RT-PCR primers to test 109J transcript truncation |
| RT_1075_GAP_R | TGGCTCAGACGCTCTTGC |  |

| **Supplementary Table 4. Plasmids used in this research study.** | | |
| --- | --- | --- |
| **Plasmid** | **Description** | **Source** |
| pK18*mobsacB* | Suicide vector (kan^R^, *lacZ*α, *sacB*) used for crossovers into the *B. bacteriovorus* genome | ^8^ |
| pMQBAD | Plasmid capable of replication within *B. bacteriovorus* and used for complementation (tdTomato, gent^R^) | This lab. Originally derived from pMQ414^9^ |
| pAKF56 | Template for *mCherry* gene | ^10^ |
| pdelta1075 | Upstream and downstream fragments of *bd1075* gene for unmarked gene deletion | This study |
| p1075_HD100c | *bd1075* from strain HD100 for complementation | This study |
| p1075_109Jc | *bd1075* from strain 109J for complementation | This study |
| p1075mCh_DXO | Full length Bd1075-mCherry fusion (double-crossover) | This study |
| p1075mCh_SXO | Full length Bd1075-mCherry fusion (single-crossover) | This study |
| p1075mCh_E302_SXO | Bd1075-mCherry fusion terminating at Bd1075 residue E302 (single-crossover) | This study |
| p1075mCh_A304_SXO | Bd1075-mCherry fusion terminating at Bd1075 residue A304 (single-crossover) | This study |
| p1075mCh_C156A_SXO | Full length Bd1075-mCherry fusion with a point mutation of C156A (single-crossover) | This study |
| p1075mCh_Y274A_SXO | Full length Bd1075-mCherry fusion with a point mutation of Y274A (single-crossover) | This study |
| pbd0064-mCeru_SXO | Bd0064-mCerulean3 (single-crossover) | This study |

| **Supplementary Table 5. Strains used in this research study.** | | |
| --- | --- | --- |
| **Strain** | **Description** | **Source** |
| *E. coli* DH5α | *E. coli* cloning strain (F- endA1 hsdR17 (rk -mk -) supE44 thi-1 recA1 gyrA (NaIr) relA1 Δ(lacIZYA-argF) U169 deoR (80dlacΔ (lacZ)M15)) | ^11^ |
| *E. coli* S17-1 | *E. coli* strain (thi, pro, hsdR-, hsdM+, recA; integrated plasmid RP4- Tc::Mu-Kn::tn) | ^12^ |
| *E. coli* S17-1: pZMR100 | *E. coli* strain containing the plasmid pZMR100 (kan^R^) | ^13^ |
| *E. coli* S17-1: pUC19 | *E. coli* strain containing the plasmid pUC19 (gent^R^) | This lab |
| *E. coli* S17-1: pK18_Bd0064mCerulean3_SXO | *E. coli* strain with pK18*mobsacB* plasmid containing Bd0064-mCerulean3 fusion (single-crossover) | This study |
| *E. coli* S17-1: lux | *E. coli* S17-1 strain (luminescent) | Gift from Dr Philip Hill |
| *B. bacteriovorus* HD100 | *B. bacteriovorus* Type strain, genome-sequenced, wild-type | ^14^ |
| *B. bacteriovorus* 109J | *B. bacteriovorus* strain, genome-sequenced, wild-type | This lab |
| *B. bacteriovorus* HD100 Δ*bd1075* | *B. bacteriovorus* containing an in-frame unmarked deletion of *bd1075* | This study |
| *B. bacteriovorus* HD100 Δ*bd1075:* p1075_HD100c | HD100 Δ*bd1075* containing the complementation plasmid p1075_HD100c | This study |
| *B. bacteriovorus* HD100 Δ*bd1075*: p1075_109Jc | HD100 Δ*bd1075* containing the complementation plasmid p1075_109Jc | This study |
| *B. bacteriovorus* HD100 Δ*bd1075*: pMQBAD | HD100 Δ*bd1075* containing the empty complementation plasmid | This study |
| *B. bacteriovorus* 109J: p1075_HD100c | 109J containing the complementation plasmid p1075_HD100c | This study |
| *B. bacteriovorus* 109J: pMQBAD | 109J containing the empty complementation plasmid | This study |
| *B. bacteriovorus* HD100 Bd1075mCh_DXO | HD100 containing a double-crossover, full length Bd1075-mCherry fusion | This study |
| *B. bacteriovorus* HD100 Bd1075mCh_SXO | HD100 containing a single-crossover, full length Bd1075-mCherry fusion | This study |
| *B.bacteriovorus* HD100 Bd1075mCh_DXO_64mCerulean3_SXO | HD100 containing both a double-crossover, full length Bd1075-mCherry fusion and a single-crossover Bd0064-mCerulean3 fusion. | This study |
| *B. bacteriovorus* HD100 Bd1075mCh_E302_SXO | HD100 containing a single-crossover Bd1075-mCherry fusion terminating at Bd1075 residue E302 | This study |
| *B. bacteriovorus* HD100 Bd1075mCh_A304_SXO | HD100 containing a single-crossover Bd1075-mCherry fusion terminating at Bd1075 residue A304 | This study |
| *B. bacteriovorus* HD100 Bd1075mCh_C156A_SXO | HD100 containing a single-crossover, full length Bd1075-mCherry fusion with the point mutation C156A | This study |
| *B. bacteriovorus* HD100 Bd1075mCh_Y274A_SXO | HD100 containing a single-crossover, full length Bd1075-mCherry fusion with the point mutation Y274A | This study |
| *B. bacteriovorus* HD100 Δ*bd1075* Bd1075mCh_SXO | HD100 Δ*bd1075* containing a single-crossover, full length Bd1075-mCherry fusion | This study |
| *B. bacteriovorus* HD100 Δ*bd1075* Bd1075mCh_E302_SXO | HD100 Δ*bd1075* containing a single-crossover Bd1075-mCherry fusion terminating at Bd1075 residue E302 | This study |
| *B. bacteriovorus* HD100 Δ*bd1075* Bd1075mCh_A304_SXO | HD100 Δ*bd1075* containing a single-crossover Bd1075-mCherry fusion terminating at Bd1075 residue A304 | This study |
| *B. bacteriovorus* HD100 Δ*bd1075* Bd1075mCh_C156A_SXO | HD100 Δ*bd1075* containing a single-crossover, full length Bd1075-mCherry fusion with the point mutation C156A | This study |
| *B. bacteriovorus* HD100 Δ*bd1075* Bd1075mCh_Y274A_SXO | HD100 Δ*bd1075* containing a single-crossover, full length Bd1075-mCherry fusion with the point mutation Y274A | This study |
| *B. bacteriovorus* HD100 Bd0064mCeru_SXO | HD100 containing a single-crossover Bd0064-mCerulean3 fusion | This study |
| *B. bacteriovorus* HD100 Δ*bd1075* Bd0064mCeru_SXO | HD100 Δ*bd1075* containing a single-crossover Bd0064-mCerulean3 fusion | This study |

**Supplementary methods**

**Reverse-transcriptase PCR to assay gene transcription during the *B. bacteriovorus* HD100 predatory cycle**
To monitor the expression of *bd1075* across the predatory cycle, RNA template was isolated from different time points during a synchronous predation of *B. bacteriovorus* HD100 on *E. coli* S17-1 using an SV Total RNA Isolation System kit (Promega) as previously described^15^. RNA quality was verified on an Agilent Bioanalyzer using an Agilent RNA 6000 Nano Kit. RT-PCR was carried out using the QIAGEN OneStep RT-PCR kit with the following thermocycling parameters: 50 °C for 30 min, 94 °C for 15 min, followed by 30 cycles of 94 °C for 1 min, 50 °C for 1 min, and 72 °C for 1 min, and a final step of 72 °C for 10 min.

***B. bacteriovorus* predation on *E. coli* in liquid culture**

The predation efficiency of *B. bacteriovorus* strains in liquid culture was measured using a luminescent prey assay developed by Lambert *et al*., 2003^16^. *B. bacteriovorus* strains were cultured in Ca/HEPES buffer at 29 °C with orbital shaking at 200 rpm for 24 h. Wild-type *B. bacteriovorus* HD100 and Δ*bd1075* predator strains were then matched by total protein content which was determined by Lowry assay^17^. Matched *B. bacteriovorus* strains were enumerated on overlay plates upon which plaques emerged after 5-7 days. Luminescent *E. coli* S17-1 was cultured in YT broth at 37 °C with orbital shaking at 200 rpm for 16 h. The stationary phase *E. coli* was then adjusted to OD_600_ 0.2 in a 1:1 mixture of Ca/HEPES and PY medium and 200 μl were aliquoted into each well of a 96-well microtiter optiplate. *E. coli* prey were enumerated on YT agar plates at 37 °C for 16 h and typically had a concentration of 10^7^ CFU ml^-1^. One millilitre samples of *B. bacteriovorus* were heat-killed by incubation at 105 °C for 5 min. To create a predator dilution series, *B. bacteriovorus* live cell volumes of 0, 1, 2, 4, 8, 16, 32 and 64 μl were aliquoted into each well containing *E. coli* prey and made up to a total of 64 μl with heat-killed predator cells. Microtiter plates were covered with a Breath-Easy® membrane and reduction in *E. coli* luminescence was measured at 30 min intervals in a BMG FluoStar microplate reader maintained at 29 °C with double orbital shaking at 200 rpm. To compare the rates of *E. coli* death by predation, the area under each luminescence curve was measured and then normalized to the maximum luminescence for each dilution. Data were analyzed in BMG LABTECH MARS data analysis software.

***B. bacteriovorus* predation on *E. coli* biofilms**

The predation efficiency of *B. bacteriovorus* strains on prey biofilms was measured using an assay adapted from Lambert & Sockett, 2013^18^ and Medina *et al.*, 2008^19^. *E. coli* S17-1 was cultured in YT broth at 37 °C with orbital shaking at 200 rpm for 16 h and then 200 μl of stationary phase *E. coli*, adjusted to OD_600_ 0.1 in fresh YT, were aliquoted into each well of a 96-well PVC microtiter plate and incubated in a 29 °C static incubator for 24 h to produce a prey biofilm. *E. coli* prey were enumerated on YT agar plates at 37 °C for 16 h and typically had a concentration of 10^7^ CFU ml^-1^. *B. bacteriovorus* WT HD100 and Δ*bd1075* were concurrently cultured at 29 °C with orbital shaking at 200 rpm for 24 h. Following complete prey lysis, *B. bacteriovorus* predators were filtered through a 0.45 μm membrane to remove residual prey. Predator strains were then matched by total protein and plaques were enumerated on double-layer overlay plates after 5-7 days. The *E. coli* biofilm-coated plate was removed from the incubator and washed carefully three times with Ca/HEPES buffer to remove residual planktonic cells. *B. bacteriovorus* 200 μl aliquots of neat (10^0^), 10^-1^, 10^-2^ and 10^-3^ dilutions in Ca/HEPES buffer were then added to the *E. coli* biofilm plate. *B. bacteriovorus* filtered through a 0.22 μm membrane served as a no-predator control. The 96-well plate containing predators and prey was incubated for a further 24 h in a static 29 °C incubator. The plate was then stained with 1% crystal violet for 15 min, washed with sterile distilled water, and de-stained with 33% acetic acid for 15 min. Plate contents were mixed carefully and then absorbance readings were acquired at OD_600_ to quantify the amount of *E. coli* biofilm remaining.

**Supplementary information references**
